# Shifted assembly and function of mSWI/SNF family subcomplexes underlie targetable dependencies in dedifferentiated endometrial carcinomas

**DOI:** 10.1101/2025.10.13.681937

**Authors:** Jessica D. St. Laurent, Grace D. Xu, Alexander W. Ying, Bengul Gokbayrak, Ajinkya Patil, Joao A. Paulo, Kasey S. Cervantes, Shary Chen, William W. Feng, Akshay Sankar, Daniel D. Samé Guerra, Jun Qi, Dana S. Neel, Jason L. Hornick, David L. Kolin, Steven P. Gygi, David G. Hunstman, Yemin Wang, Cigall Kadoch

**Author notes:** These authors contributed equally (Grace D. Xu, Alexander W. Ying, Yemin Wang).

## Abstract

The mammalian SWI/SNF (mSWI/SNF) family of chromatin remodelers govern cell type-specific chromatin accessibility and gene expression and assemble as three distinct complexes: canonical BAF (cBAF), Polybromo-associated BAF (PBAF), and non-canonical BAF (ncBAF). ARID1A and ARID1B are paralog subunits that specifically nucleate the assembly of cBAF complexes and are frequently co-mutated in highly aggressive dedifferentiated/undifferentiated endometrial carcinomas (DDEC/UECs). Here, in cellular models and primary human tumors, we find that ARID1A/B deficiency-mediated cBAF loss results in increased ncBAF and PBAF biochemical abundance and chromatin-level functions to maintain the DDEC oncogenic state. Further, treatment with clinical-grade SMARCA4/2 ATPase inhibitors markedly attenuates DDEC cell proliferation and tumor growth in vivo and synergizes with carboplatin-based chemotherapy to extend survival. These findings reveal the oncogenic contributions of shifted mSWI/SNF family complex stoichiometry and resulting gene regulatory dysregulation and suggest therapeutic utility of mSWI/SNF small molecule inhibitors in DDEC/UEC and other cBAF-disrupted cancer types.

## Introduction

Genome-wide sequencing efforts performed over the past decade have revealed extensive involvement of chromatin regulatory proteins and protein complexes in human disease^1,2^. Of note, the genes encoding the mammalian SWI/SNF (mSWI/SNF) family of chromatin remodeling complexes are among the most frequently perturbed entities in human cancer and in neurodevelopmental disorders, including several disease pathologies driven uniformly by mSWI/SNF subunit compromise^3–5^. Rare cancers such as malignant rhabdoid tumor, synovial sarcoma, small cell carcinoma of the ovary, hypercalcemic type (SCCOHT), as well as intellectual disability syndromes such as Coffin-Siris syndrome, are considered to be caused by (and in some cases, also diagnosed by) specific mSWI/SNF perturbations^6–12^, providing unique windows into the mechanisms by which these complexes control chromatin architecture and regulate gene expression. Studies by our group and others have leveraged such settings to inform subunit-specific contributions to mSWI/SNF activity, delineate subunit biochemical assembly and structural relationships, and identify unique vulnerabilities that can be accompanied by novel therapeutic approaches^13–21^.

Recently, targeted sequencing and immunohistochemical characterization of a uniquely aggressive form of endometrial cancer, dedifferentiated/undifferentiated endometrial carcinoma (DDEC/UEC), revealed a high prevalence of mSWI/SNF perturbations in over 70% of cases^22–24^. DDEC is an epithelial malignancy comprising a UEC that arises in the background of a well or poorly differentiated endometrial carcinoma^23^. Among mSWI/SNF mutations, concomitant mutation of ARID1A and ARID1B subunits is most common (Coatham et al. and DFCI cohorts^25^), with SMARCA4 and SMARCB1 as other frequently disrupted subunit genes (**Fig. 1A, Extended Data Fig. 1A-E, Supplementary Table 1**) ^25^. Loss of both ARID1A and ARID1B expression is often observed in the undifferentiated component whereas the corresponding differentiated component shows intact expression of ARID1B^22,25^. Mutations in *ARID1A* are also common in well-differentiated endometrial carcinomas, occurring in approximately 40% of cases, while *SMARCA4/2* mutations are infrequent^26^. Furthermore, DDEC cases can exhibit mutations in *POLE* and *TP53* (**Extended Data Fig. S1A-C**) ^24,27,28^. Diagnosis of UEC is made based on histologic tumor appearance and supported by the loss of epithelial linage markers including PAX8, cytokeratins, estrogen receptor, progesterone receptor and E-cadherin^29^. Of note, compared to mSWI/SNF-intact DDEC cases, ARID1A/B-dual mutated cases result in significantly shorter overall survival (∼4 months versus ∼36 months) and increased resistance to conventional chemotherapy with advanced stage disease^22,24,25^. Taken together, these studies underscore that concomitant ARID1A/B loss defines a highly aggressive group of undifferentiated endometrial cancers characterized by rapid disease progression and insensitivity to conventional platinum and taxane-based chemotherapy. Here we define the biochemical and chromatin regulatory mechanisms underlying ARID1A/B dual-deficient DDEC/UEC cancers, and inform new potential therapeutic approaches.

**Figure 1.**
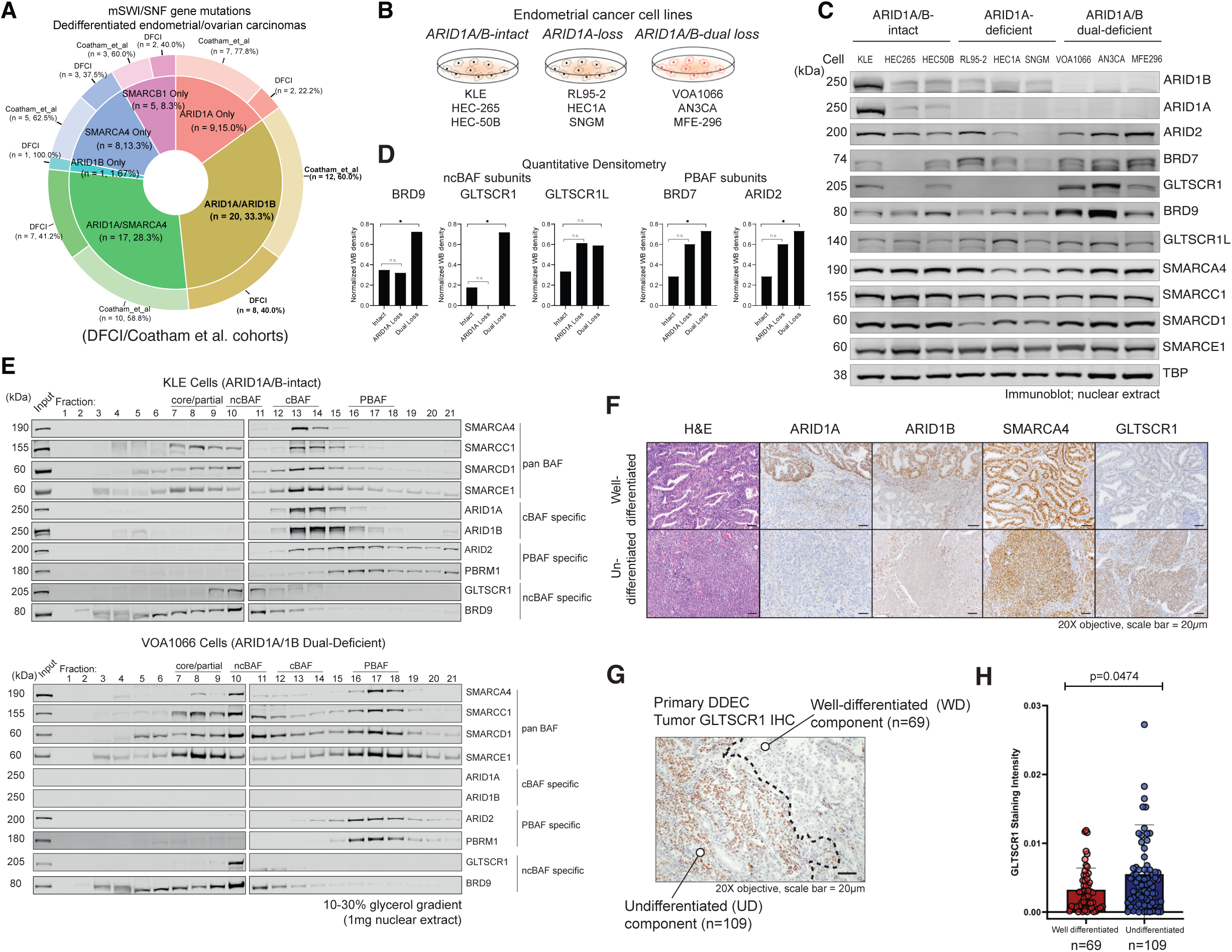
ARID1A/ARID1B dual-deficient endometrial tumors exhibit cBAF loss and increased abundance of ncBAF and PBAF complex components. A. Pie chart reflecting targeted sequencing of mSWI/SNF-mutant dedifferentiated endometrial, and ovarian carcinomas (DFCI and Coatham et al., 2016 cohorts) indicating mutational frequencies of mSWI/SNF genes. Dual ARID1A and ARID1B mutations are present in ∼33% of DDEC cases. B. Schematic highlighting n=3 ARID1A/B-intact, ARID1A-loss and ARID1A/B-dual-loss endometrioid cancer cell lines used in this study. C. Immunoblot for selected mSWI/SNF subunits spanning all 3 complex types performed on nuclear extracts isolated from ARID1A/B-intact, ARID1A-loss, and ARID1A/B dual-deficient endometrial cell lines. D. Quantitative densitometry analyses performed on immunoblots for ncBAF subunits (BRD9, GLTSCR1) and PBAF subunits (BRD7, ARID2) in ARID1A/B-intact, ARID1A-loss, and ARID1A/B dual-deficient endometrial cell lines, n=3 per condition, *p<0.05. E. Density sedimentation studies using 10-30% glycerol gradients performed on nuclear extracts isolated from KLE and VOA1066 cell lines and immunoblotted for selected mSWI/SNF subunits, n=3 (one replicate shown). F. H&E and immunostaining for ARID1A, ARID1B, SMARCA4 and GLTSCR1 performed on representative DDEC cases, showing well-differentiated and undifferentiated compartments, n=75. G. Representative photomicrograph of DDEC primary tumor GLTSCR1 IHC, indicating well-differentiated and undifferentiated compartments; number of cases used in analyses are indicated. H. Bar graph quantifying GLTSCR1 staining intensity in DDEC well-differentiated and undifferentiated compartments. p-values determined with two-tailed Students t-test for D, H, Bars represent S.E.M.

## Results

### Dual ARID1A/B loss increases residual mSWI/SNF protein abundance

We first curated tumor-derived cell lines bearing intact expression of full-length ARID1A and ARID1B, those with loss of expression of ARID1A, and those characterized by dual-loss of ARID1A and ARID1B, isolated nuclear material and performed immunoblot analyses for mSWI/SNF family subunits (**Fig. 1B-C, Extended Data Fig. 1F, Supplementary Table 2**). Notably, among all endometrial lines, >55% are scored to contain loss-of-function mutations in *ARID1A*, consistent with high percentages found in human tumors^25,30,31^ (**Fig. 1C**, **Extended Data Fig. 1F**). The VOA1066 cell line lacks expression of ARID1A and ARID1B, as shown using immunoblot and quantitative proteomics (**Extended Data Fig. 1G-E, Supplementary Table 3**). In DDEC tumors, while cBAF-specific ARID1A/B subunits were entirely absent, we found that ncBAF-specific subunits, BRD9, GLTSCR1, and GLTSCR1L as well as PBAF subunits, ARID2 and BRD7, were increased in protein-level abundance in these cell lines (**Fig. 1C-D**). To further examine this, we subjected nuclear extracts of an mSWI/SNF-intact cell line, KLE, and the ARID1A/B-dual deficient cell line, VOA1066, to density sedimentation analyses using 10-30% glycerol gradients (**Fig. 1E**). Notably, subunits including SMARCC1 and SMARCD1, core module subunits that can nucleate both ncBAF and cBAF complexes, exhibited clear shifts toward the lower (fractions 8-11) and upper (fractions 16-18) molecular weight fractions in the VOA1066 cell line relative to the KLE cell line (**Fig.1E**). Further, we noted increases in BRD9 and GLTSCR1/1L ncBAF subunit intensities (relative to total input) in fractions 6-11 and PBAF-associated PBRM1 intensities in the dual-deficient VOA1066 cell line relative to the KLE mSWI/SNF-intact cell line (**Fig. 1E**). Finally, we performed comprehensive immunohistochemical profiling on a collection of n=109 primary human DDEC tumors bearing both undifferentiated (UD) (n=109) and well-differentiated (WD) (n=69) regions (**Fig.1F-G, Extended Data Fig. 1I-E**). Immunostaining of GLTSCR1, a nucleating member of ncBAF complexes, was significantly elevated in de-differentiated tumor compartments, with similar intensities of the ATPase subunit, SMARCA4, across both UD and WD compartments (**Fig.1H, Extended Data Fig. 1K**). Thus, remaining mSWI/SNF subcomplexes exhibit altered subunit protein-level abundance in both ARID1A/B-deficient cellular models and primary human tumors.

### ARID1A restoration results in mSWI/SNF rebalance

To determine whether the above observations in cell lines and primary tumors were specifically due to the loss of the ARID1A/B subunits, we next generated a system for the restoration of full-length WT ARID1A in DDEC cell lines (**Fig. 2A**). Indeed, ARID1A rescue into both VOA1066 and AN3CA cells resulted in reduced nuclear protein levels of ncBAF components (BRD9 in both cell lines and GLTSCR1 in VOA1066 cells most prominently) as well as PBAF (ARID2) components in both cell lines (**Fig. 2B**). Importantly, we observed a clear redistribution of ncBAF-cBAF shared core module subunits, such as SMARCC1 and SMARCD1, as well as ATPases SMARCA4 and SMARCA2 using density sedimentation (**Fig. 2C, Extended Data Fig. 2A**). Quantitative tandem mass-tag (TMT) proteomics performed on total cell lysates confirmed these findings, revealing significant changes in subunit protein abundance of ncBAF and PBAF subunits and restoration of cBAF abundance following ARID1A rescue (**Fig. 2D-E, Extended Data Fig. 2B-E, Supplementary Table 4**). Importantly, shifted complex assembly was accompanied by significant reductions in proliferation and altered morphology suggestive of endometrial differentiation in DDEC cell lines in culture (**Fig. 2F, Extended Data Fig. 2E-E**).

**Figure 2.**
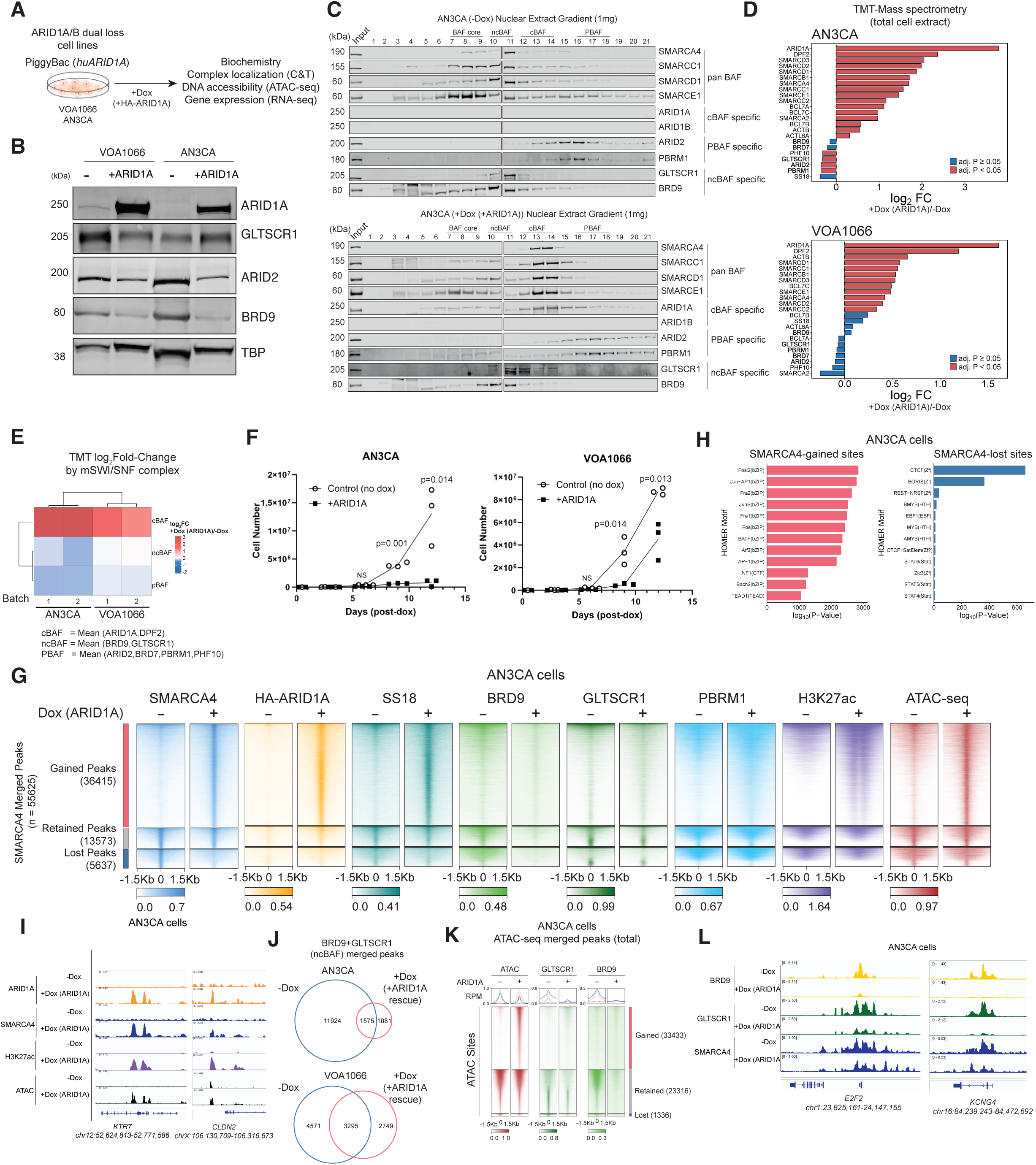
cBAF reassembly controls genome-wide redistribution of mSWI/SNF family complexes in DDEC cells. A. Schematic for overexpression of HA-tagged full-length ARID1A in AN3CA and VOA1066 cell lines using an inducible piggyBac system. B. Immunoblot for mSWI/SNF complex components spanning cBAF (ARID1A), PBAF (ARID2) and ncBAF (GLTSCR1, BRD9), along with TBP as loading control in both AN3CA and VOA1066 cell line systems in (-) and (+) Dox conditions, n=3 (one replicate shown). C. Density sedimentation analyses performed on AN3CA ARID1A/B dual-deficient cells in -Dox (control) and +Dox (ARID1A rescue) conditions using 10-30% glycerol gradients. Immunoblots for selected mSWI/SNF subunits are shown, n=3 (one replicate shown). D. Bar graphs depicting Log2FC changes in whole-cell mSWI/SNF subunit abundance in AN3CA and VOA1066 cells in −/+ Dox (ARID1A rescue) conditions defined by TMT mass-spectrometry (n=4 replicates per condition); significance scores are indicated in the legend; ncBAF- and PBAF-specific subunits are indicated in bold. E. Summed log-fold changes for peptides corresponding to cBAF, PBAF, and ncBAF subunits in VOA1066 and AN3CA cell lines upon ARID1A rescue (+Dox/-Dox logFC) shown as heatmap. F. Cell proliferation experiments in AN3CA and VOA1066 DDEC cell lines treated with either no Dox or Dox-mediated induction of ARID1A expression and number of live cells were measured on an automatic cell counter. n=5 experimental replicates; data presented are as a mean +/− SD. G. Heatmaps reflecting ChIP-seq in AN3CA cells for HA (ARID1A), SMARCA4, SS18, BRD9, GLTSCR1, PBRM1 and H3K27Ac (each normalized to input), along with ATAC-seq. Gained, lost and retained peak sets are indicated. H. HOMER motif analyses performed over gained and lost SMARCA4 sites in AN3CA;+Dox (ARID1A rescue), -Dox (no rescue control),p-values calculated using HOMER algorithm. I. Representative ChIP-seq tracks over the KRT7, and CLDN2 loci in AN3CA cells. J. Venn diagrams depicting BRD9-GLTSCR1 (ncBAF) merged peaks in −/+ Dox (ARID1A rescue) conditions in AN3CA and VOA1066 cell lines. K. Heatmaps depicting ATAC-seq signal along with BRD9 and GLTSCR1 ChIP-seq signal over gained, lost and retained ATAC-seq peaks in AN3CA cells. L. Representative ChIP-seq tracks for BRD9, GLTSCR1, and SMARCA4 over the E2F2 and KCNG4 loci in AN3CA cells.

We next sought to determine the chromatin-level changes in mSWI/SNF complex occupancy following ARID1A rescue in DDEC cell lines (**Fig. 2A**). We profiled mSWI/SNF complexes marked by SMARCA4 (present in all mSWI/SNF subcomplexes), and found that ARID1A restoration led to a large number of gained sites genome-wide (n=36415), as well as a smaller number of sites with retained occupancy (n=13573), and lost or reduced occupancy (n=5637) in AN3CA cells (**Fig. 2G, Extended Data Fig. 2H**). Sites with gained mSWI/SNF complex targeting corresponded largely to restored cBAF complex occupancy as exhibited by overlap with ARID1A and SS18, as well as increases in the H3K27Ac mark associated with active sites ^21,32^ and chromatin accessibility as profiled via ATAC-seq (**Fig. 2G**). HOMER motif enrichment and distance-to-TSS analyses revealed that SMARCA4-gained sites were largely TSS-distal, consistent with rescue of cBAF assembly and localization to distal enhancer sites^14,17^ with gained cBAF-associated motifs enriched for Fos, Jun, AP-1, and ATF3 bZIP transcription factors (TFs)^20,33^ (**Fig. 2H, Extended Data Fig. 2I**). Similar results were obtained in the VOA1066 cell line (**Extended Data Fig. 2H-E**). Sites with gained mSWI/SNF occupancy were observed over loci such as *KRT7 and CLDN2* (**Fig. 2I**). Further, in both AN3CA and VOA1066 cell lines, accessibility was gained across a large number of sites upon ARID1A rescue, corresponding to increases in cBAF complex targeting to distal regions (**Fig. 2G, Extended Data Fig. 2K-E**). These data suggest a role for ARID1A-containing cBAF complexes in restoring open chromatin architecture and activating genes such as lineage markers in DDEC.

In contrast, however, we noted that sites that exhibited reductions in SMARCA4 occupancy, as well as those with retained occupancy following ARID1A restoration were those sites showing strong reductions in ncBAF (BRD9 and GLTSCR1) occupancy (**Fig. 2G, J**). Intriguingly, these sites were not accompanied by significant changes in accessibility (**Fig. 2G, K, Extended Data Fig. 2M**), consistent with prior studies indicating a lesser role for ncBAF complexes in chromatin accessibility generation relative to cBAF^17,18^. Correspondingly, SMARCA4-marked sites lost upon ARID1A restoration were promoter-proximal and exhibited strong enrichment for CTCF and BORIS (CTCF-like) Zf TF motifs known to be targeted by ncBAF complexes^17^ (**Fig. 2H, Extended Data Fig. 2J**). We found that gained cBAF peaks following ARID1A restoration replaced between 35-45% of ncBAF peaks (**Extended Data Fig. 2N**). Further, we identified that upon ARID1A rescue, ncBAF occupancy was substantially reduced, especially over sites of both retained and lost chromatin accessibility (**Fig. 2J-K, Extended Data Fig. 2M**). This is exemplified over the *E2F2* and *KCNG4* loci (**Fig. 2L**). Finally, we examined the PBAF-specific subunit, PBRM1, finding that PBAF complexes mainly localized to retained sites in either empty or ARID1A rescue conditions, albeit with minimal changes to occupancy levels over promoters (**Extended Data Fig. 2O-E**). These data demonstrate marked changes in both protein-level abundance and genomic binding and activity of mSWI/SNF family complexes, specifically ncBAF and PBAF complexes, that are controlled by cBAF assembly in DDEC cells.

### ARID1A rescue induces differentiation and attenuates oncogenicity

We next sought to define the impact of ARID1A restoration on gene expression in DDEC. We identified both gains and losses in gene expression, with ∼4-5-fold more genes upregulated than downregulated, aligned with chromatin accessibility changes observed (**Fig. 3A, Extended Data Fig. 3A** and **Fig. 2G, Extended Data Fig. 2L**). Across both DDEC cell lines, upregulated genes included those involved in hormone response (estrogen and androgen), cell adhesion (EMT, apical junction), inflammatory response, IFNalpha/gamma response, as well as apoptosis (**Fig. 3B, Extended Data Fig. 3B-E**). Consistent with the antiproliferative impact of ARID1A restoration (**Fig. 2F, Extended Data Fig. 2E-E**), gene expression pathways most strongly downregulated included those of G2M checkpoint, MYC and E2F targets and mTORC signaling (**Fig. 3B, Extended Data Fig. 3B-E**). Concordantly upregulated genes in both DDEC cell types included *WNT6*, *PRKCA*, *SOX13* and *PRICKLE1* and downregulated genes such as *MYC*, *DDN*, and *GLI1* and *NR4A2* (**Fig. 3A-B, Extended Data Fig. 3E-E**). Integrating these data with TMT-MS, we identified proteins associated with cell-cell adhesion, tissue-specific cell differentiation, and apoptosis/cell death as upregulated, while those involved in cell cycle, DNA repair and DNA replication as downregulated at both the RNA and protein levels (**Fig. 3C**).

**Figure 3.**
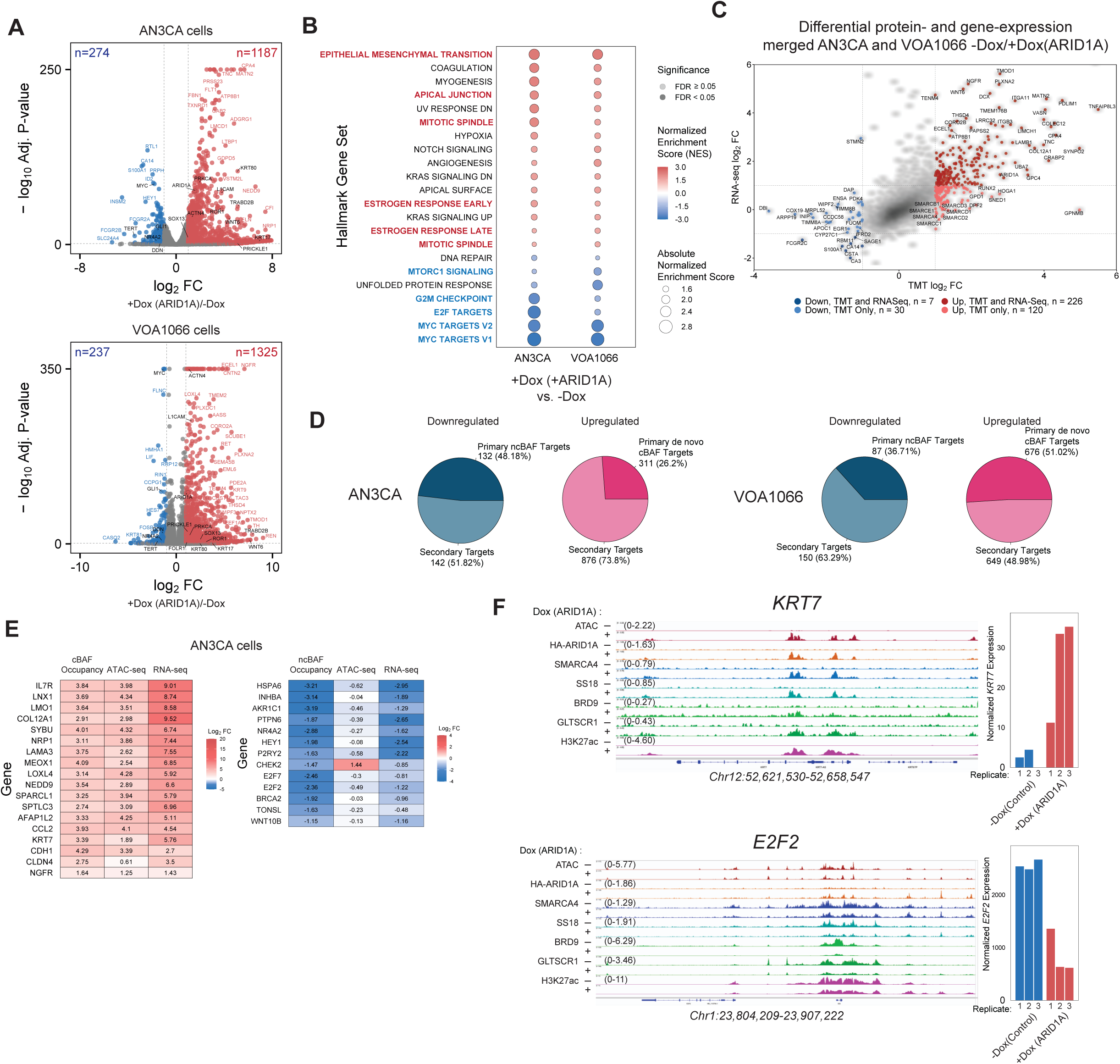
ARID1A/ARID1B-mediated mSWI/SNF complex reshuffling underlies altered gene regulation in DDEC cell lines and primary tumors. A. Volcano plots showing up- (red) and down- (blue) regulated genes following 72 hours of +Dox (ARID1A restoration) in AN3CA and VOA1066 DDEC cell lines. Adjusted p-values calculated from DESeq2. B. Hallmark GSEA showing ARID1A rescue-mediated gene expression pathway changes in AN3CA and VOA1066 cells. Genes labeled in black indicate concordantly up- and down-regulated genes. NES, normalized enrichment score. C. Scatter plot depicting significantly changed proteins (x-axis, TMT mass-spec) and RNA levels (RNA-seq) following ARID1A rescue in AN3CA and VOA1066 cells (merged datasets). D. Pie charts reflecting downregulated genes that are primary targets (defined as BRD9/GLTSCR1 co-targeted genes) and secondary targets (not ncBAF-targeted) and upregulated genes that are targeted by cBAF (defined as HA/ARID1A co-targeted site) upon rescue of ARID1A in AN3CA and VOA1066 cells. E. (Left) Log2FC of cBAF occupancy, ATAC-seq, and RNA-seq signals upon ARID1A rescue; (right) Log2FC of ncBAF occupancy, ATAC-seq, and RNA-seq. F. Representative ChIP-seq and ATAC-seq tracks at the KRT7 and E2F2 loci with corresponding bar graphs depicting gene expression (RNA-seq).

We next sought to define putative direct target functions of cBAF loss and ncBAF/PBAF complex gains and their shifted targeting genome-wide. Indeed, among ARID1A-mediated downregulated genes defined by RNA-seq experiments, ∼48% and ∼37% were genes near sites of lost ncBAF (BRD9-/ GLTSCR1-shared sites) targeting in AN3CA and VOA1066 cells, respectively (**Fig. 3D**). Approximately 26% and 51% of upregulated genes were those nearby sites of increased cBAF chromatin occupancy and accessibility (HA-ARID1A-/SMARCA4-shared sites) in AN3CA and VOA1066 cells, respectively (**Fig. 3D**). As examples, we defined increased cBAF targeting, accessibility and gene expression at the *IL7R*, *NRP1*, and *KRT7* loci and concomitant decreased ncBAF targeting and gene expression, coupled with more minor changes in accessibility over the *E2F2*, *WNT10B* and *INHBA* genes in AN3CA cells (**Fig. 3E, Extended Data Fig. 3E-E**). This is exemplified at the *KRT7* and *E2F2* loci, respectively (**Fig. 3F**, **Extended Data Fig. 3H**) and together suggests DDEC cell endometrial redifferentiation and reactivation of pathways such as hormone response. Genes with reduced expression were nearest sites of reduced ncBAF complex occupancy and accessibility and associated with tumor cell invasion, EMT, DNA repair and cell proliferation (**Fig. 3D-F, Extended Data Fig. 3F-E**).

### ARID1A/B have distinct and shared gene regulatory functions

Given that DDEC tumors exhibit concomitant loss of ARID1A and ARID1B paralog subunits of cBAF complexes, we next sought to define the impact of restoration of each paralog in isolation and together (**Fig. 4A-B, Extended Data Fig. 4A-E**). Following confirmation of protein expression of HA-tagged ARID1A, ARID1B, and both paralogs together in AN3CA cells, we performed RNA-seq and ATAC-seq studies to define the shared and disparate gene regulatory and chromatin accessibility profiles, respectively. Intriguingly, over two thirds (67.3%) of ARID1A-mediated upregulated genes were also upregulated by restoration of ARID1B; ARID1B, however, resulted in more upregulated genes than ARID1A (n=1720 versus n=1187 for ARID1A rescue) (**Fig. 4C-D**). For downregulated genes, ARID1B also induced more downregulated genes relative to ARID1A (n=1136 versus n=274 for ARID1A) (**Extended Data Fig. 4C**). Notably, 589 genes were uniquely upregulated and 622 genes uniquely downregulated only with the rescue of both ARID1A and ARID1B rather than either alone (**Fig. 4D, Extended Data Fig. 4C**). Clustering analyses revealed ‘synergistic genes’, including those such as *DUSP10*, *RARS2*, and *NFE2L* as upregulated genes and *EBP*, *GEMIN4*, *NONO* and *SP1* as downregulated genes (**Fig. 4E)**, which also exhibited concordant changes in accessibility as assessed by ATAC-seq and corresponded to adipogenesis and FLT3 signaling pathways (upregulated) and JAK2 targets, fatty acid metabolism, and endocrine therapy resistance (downregulated) (**Fig. 4E, Extended Data Fig. 4D**). The gene regulatory impact of ARID1A or ARID1B or both paralogs (majority of genes) included restoration of estradiol response, silencing of proliferative pathways and induction of senescence pathways, upregulation of SMARCA4 target genes, and downregulation of G2/M and DNA replication signatures (**Extended Data Fig. 4D**). Of note, specific TF target genes (i.e. STAT3 targets, HOXA5 targets, FOXA2 targets, and AP1 pathway targets were only upregulated significantly with restoration of ARID1B or both paralogs, but not ARID1A alone (**Extended Data Fig. 4D**). Notably, sites of increased accessibility were largely conserved between single ARID1 paralogs and their concomitant expression, with only 468 sites among >54,000 total sites showing increased accessibility only in the dual rescue condition (+ARID1A/+ARID1B) and largely mapped to synergistic genes (**Fig. 4D-E**).

**Figure 4.**
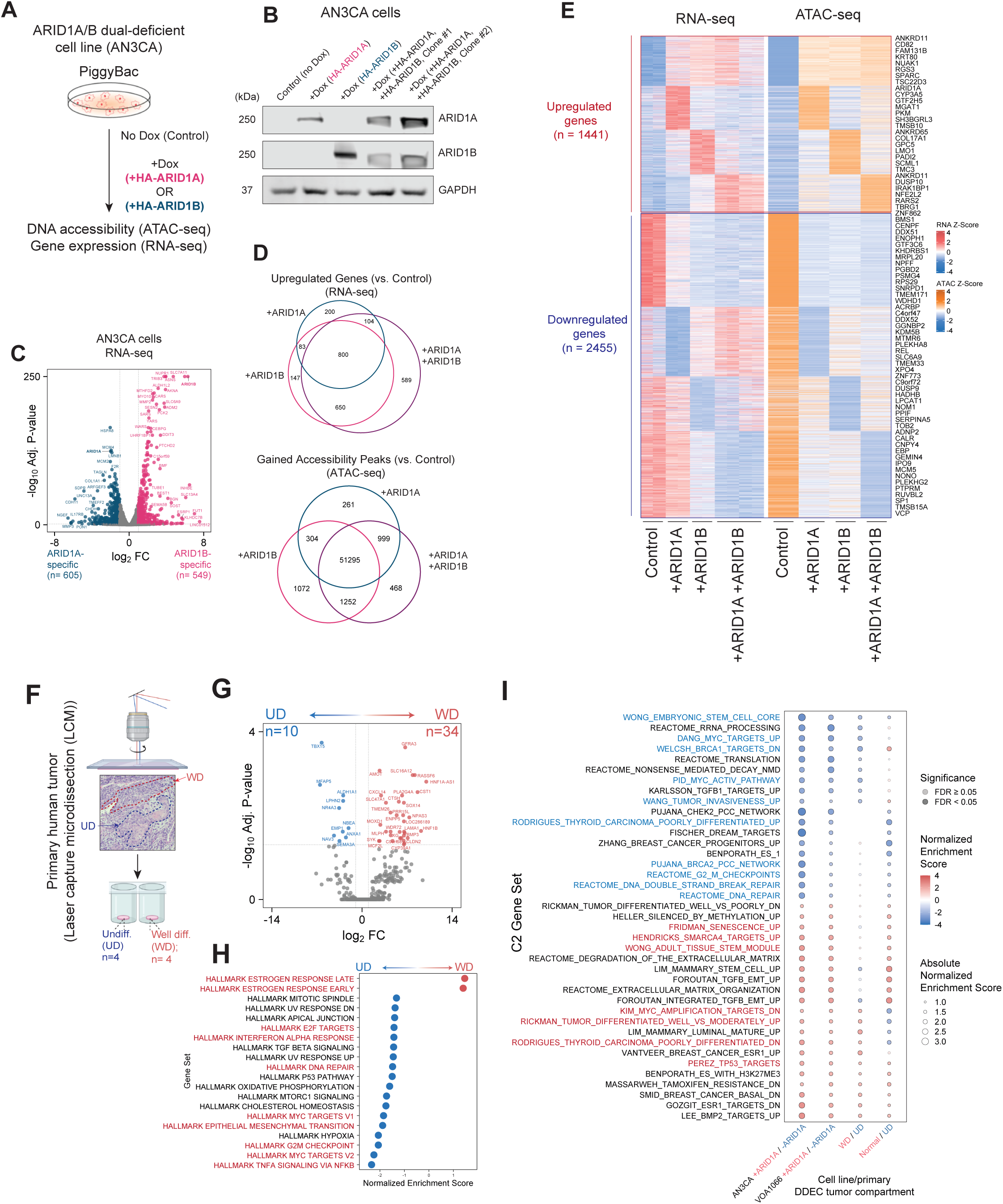
ARID1A and ARID1B paralogs mediate shared and distinct gene regulatory pathways in human DDEC cell lines and primary tumors. A. Schematic of experiments in AN3CA cells with rescue of either HA-tagged ARID1A or ARID1B (+Dox). B. Immunoblot for ARID1A, ARID1B and GAPDH performed on whole cell lysates from AN3CA cells in indicated conditions, n=3 (one replicate shown). C. Volcano plot indicating ARID1A- and ARID1B-specific upregulated genes. D. Venn diagrams indicating overlap of upregulated genes (left) and sites with increased accessibility (right) following ARID1A, ARID1B or ARID1A/B dual rescue (+Dox). E. Heatmaps depicting k-means clustered sites with ATAC-seq signal and RNA-seq signal shown as z-scores. Representative genes are labeled. F. Schematic depicting laser capture microdissection (LCM) performed on undifferentiated (UD) and well-differentiated (WD) sections of primary human endometrial tumors. Created in BioRender. G. Genes significantly enriched in UD and WD primary tumor components. H. Hallmark GSEA performed on UD- and WD-specific genes. Selected pathways upregulated (red) and downregulated (blue) are highlighted. I. GSEA (C2 gene set) performed on AN3CA and VOA1066 cells with +Dox ARID1A rescue (relative to control), WD compared to UD, and normal endometrium compared to UD. Adjusted p-values calculated from DESeq2 in (C) and (G). Selected pathways upregulated (red) and downregulated (blue) are highlighted.

Finally, we next isolated n= 4 primary human ARID1A/B dual-deficient DDEC tumors and performed laser capture microdissection (LCM) to isolate RNA from well-differentiated (WD) and undifferentiated (UD) compartments of DDEC tumors and normal tissue (**Fig. 4F, Supplementary Table 5**). We identified genes such as *ALDH1A1*, *TBX15*, and *MFAP5* as significantly upregulated genes in UD regions of all tumors, with genes such as *SOX14*, *CLDN2*, *CXCL14* and transporter genes *SLC16A12* and *SLC47A1* as upregulated in the WD regions (**Fig. 4G**). We identified that genes corresponding to hormone response (estrogen response early/late genes) as significantly upregulated in the WD regions, while genes corresponding to DNA repair, MYC targets, EMT, cell cycle and IFNa/TNFa signaling were upregulated in the UD compartments (**Fig. 4G-H**). Importantly, comparing gene sets of genes differentially expressed in the UD and WD components of primary tumors, those in normal endometrium, and those in cell line models, VOA1066 and AN3CA, we identified several clusters of genes that exhibited similar gene expression profiles and that were impacted by ARID1A rescue (**Fig. 4I, Extended Data Fig. 4E-E**), underscoring the utility of DDEC cellular models in mirroring the expression of primary tumors.

### ncBAF/ PBAF disruption attenuates DDEC phenotypes

To determine the proliferative impact of mSWI/SNF complex disruption, we generated AN3CA and VOA1066 cell lines with inducible shRNAs targeting either BRD9 (ncBAF) or ARID2 (PBAF) and assessed cell growth using colony formation assays at 6 days post induction (**Fig. 5A**). Indeed, suppression of either BRD9 and ARID2 attenuated both AN3CA and VOA1066 cell line proliferation (**Fig. 5A-B, Extended Data Fig. 5A-E).** Further, CRISPR knockout (KO) studies of ARID2, GLTSCR1, GLTSCR1L, SMARCA4 (BRG1) and SMARCA2 (BRM) indicated that each resulted in suppression of AN3CA proliferation (**Extended Data Fig. 5D-E**).

**Figure 5.**
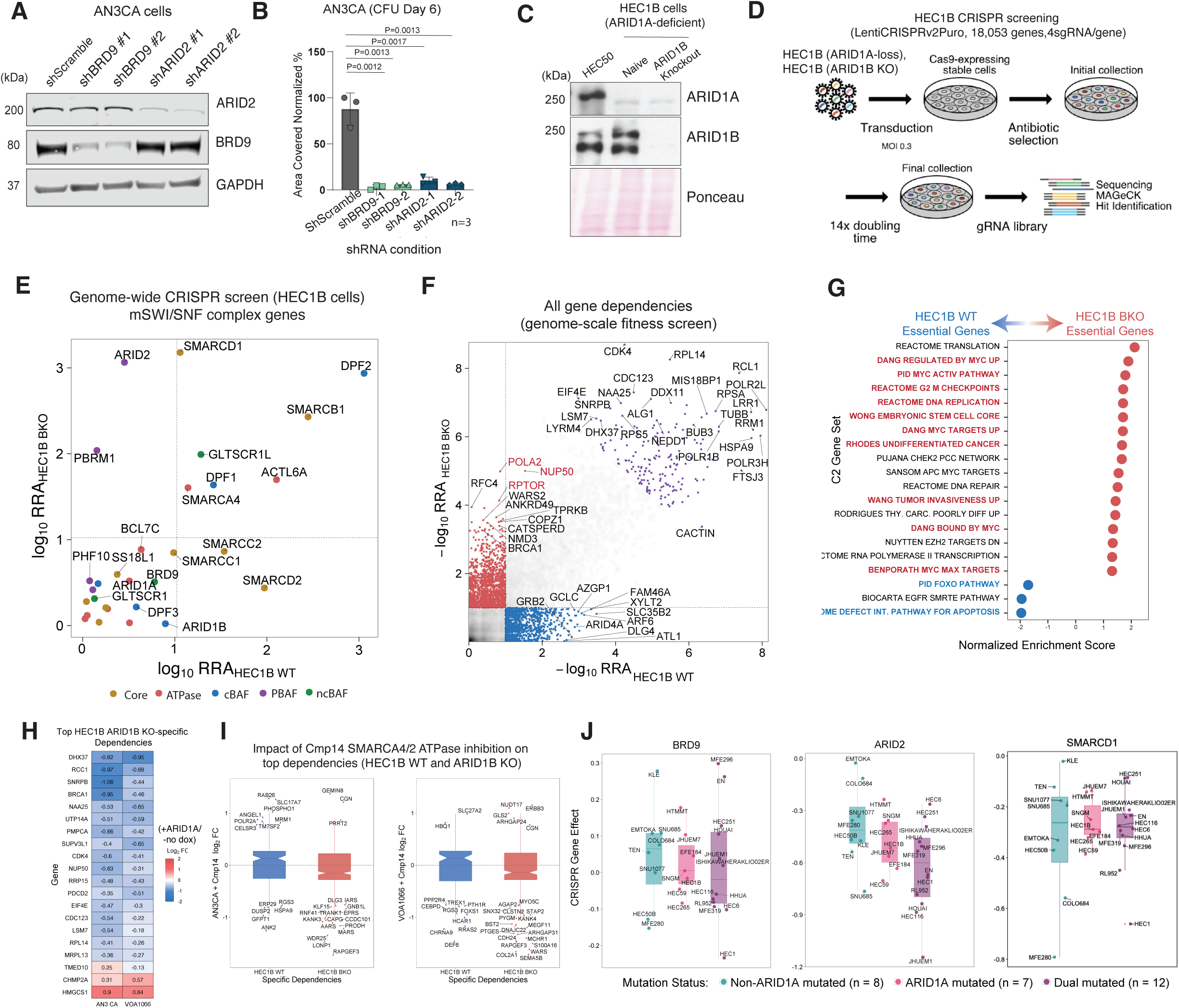
Dual ARID1 paralog-deficient DDEC cells are sensitive to genetic disruption of ncBAF and PBAF components. A. Immunoblot performed on whole-cell extract from AN3CA cells treated with indicated shRNAs targeting BRD9 or ARID2. B. Bar graph depicting AUC values in indicated conditions in AN3CA colony formation assays. C. Immunoblot performed on whole-cell extracts from HEC-50B cells and HEC1B cells treated with either non-targeting sgRNA or sgRNA targeting ARID1B, n=3 (one replicate shown). D. Schematic for CRISPR screening experiments in HEC1B cells (naive and ARID1B KO conditions). E. Scatter plot indicating CRISPR screening score (RRA) in naive and ARID1B KO HEC1B cells. F. Dependencies specific to HEC1B WT (Blue), HEC1B ARID1B KO (red) and both cell conditions (purple) displayed in -log10 RRA; significant top hits in each are indicated. G. GSEA performed on HEC1B WT cells and in HEC1B ARID1B KO conditions. H. Impact of ARID1A rescue in AN3CA and VOA1066 cell lines on top HEC1B ARID1B KO-specific dependency genes displated as Log2FC. I. Dependencies in HEC1B and ARID1B KO HEC1B cells overlapping with impact by mSWI/SNF ATPase inhibition (Cmp14 treatment), n=3 biologic replicates. J. Box and whisker plots indicating mean dependency scores (CRISPR gene effect) for BRD9, ARID2, and SMARCD1 across DepMap (Broad Institute) in endometrial cell lines with mutation conditions indicated in the legend. In (I) and (J), middle represents the median, 25%-75% and whisker + /-1.5 times the interquartile range. p-values determined with two-tailed Students t-test for (B). Error bars represent mean +/− S.E.M.

To define whether the dual loss of ARID1A/1B loss sensitized DDEC cell lines to the depletion of mSWI/SNF components, specifically, those corresponding to residual cBAF and PBAF complexes, we we used the ARID1A-deficient HEC1B endometrial cancer cell line in which we either treated with control CRISPR gRNAs or those targeting ARID1B, generating a dual ARID1A/ARID1B-deficient HEC1B cell line (**Fig. 5C**). Using genome-wide CRISPR/Cas9 screens, we defined genes that selectively impacted the proliferation of endometrial cancer cells lacking both ARID1 cBAF paralogs (**Fig. 5D, Extended Data Fig. 5F-E, Supplementary Table 6**). Notably, we identified increased dependencies on PBAF components PBRM1 and ARID2, as well as ncBAF components GLTSCR1L and SMARCD1 in ARID1A/ARID1B dual deficient HEC1B cells which we validated (**Fig. 5E, Extended Data Fig. 5H-E**). Interestingly, depletion of BRD9 resulted in proliferative attenuation of both HEC1B parental control (ARID1A-deficient) and HEC1B-ARID1B KO cells (**Extended Data Fig. 5J-E**), consistent with results of the CRISPR screen (**Fig. 5E**). Hits that scored as selective dependencies in the dual ARID1 paralog-deficient setting (**Fig. 5F**) included *POLA2*, *RAPTOR*, and *NUP50*, corresponding to pathways such as MYC targets, G2/M checkpoint, DNA replication and undifferentiated cancer signatures (**Fig. 5F-G**). Beyond PBAF and ncBAF components, several top hits included those with accompanying therapeutic opportunities, such as BRCA1 (BRCA1 inhibitors), MAP3K2 (pazopanib), SLC2A1 (WZB117), among others. To dissect top-scoring hits and define which dependencies may result from heightened ncBAF and PBAF mediated gene regulation, we identified the top 20 dependencies specific to the HEC1B ARID1B KO setting and defined their changes in gene expression in AN3CA and VOA1066 DDEC cell lines upon ARID1A restoration (**Fig. 5H, Extended Data Fig. 5K**). Further, evaluating the overlap between top dependencies in HEC1B WT and ARID1B KO settings and those genes in DDEC cell lines AN3CA and VOA1066 impacted by SMARCA4/2 ATPase inhibition using Cmp14^34^ in AN3CA and VOA1066 cell lines, we identified several genes including *KANK3*, *KANK4*, *KLF15*, *MYO5C* and *STAP2* (**Fig. 5I**). Finally, across endometrial cell lines of different ARIDA/B status, in line with our experimental findings, cell lines with either ARID1A-only or dual ARID1A/ARID1B losses exhibited increased reliance on the ncBAF components, BRD9 and SMARCD1, and the PBAF component ARID2 (**Fig. 5J**).

### Pharmacologic targeting of residual mSWI/SNF complexes

We next aimed to define the utility of small molecule inhibitors of mSWI/SNF complex ATPase activity, including Cmp14^34^ as well as dBRD9-A, a small molecule degrader of the BRD9 component of ncBAF^17,35^ (**Fig. 6A**). Treatment with either agent resulted in marked attenuation in cell proliferation of ARID1A/B dual-deficient DDEC cells in culture (**Fig. 6B, Extended Data Fig. 6A**), with Cmp14 resulting in most complete attenuation (**Fig. 6B, Extended Data Fig. 6A**). We next performed RNA-seq experiments in both VOA1066 and AN3CA cells treated with either Cmp14 or dBRD9-A for 72 hours (**Extended Data Fig. 6B-E**). Genes downregulated by both agents included those corresponding to stem cell signatures, SMARCA4 target genes, EMT, and MYC targets, tumor invasiveness and TGFB1 signatures (**Fig 6C-D, Extended Data Fig. 6D-E**). Upregulated gene sets included EZH2 targets, estrogen response, E2F targets, and cholesterol biosynthesis (**Fig. 6D, Extended Data Fig. 6D-E**). Intriguingly, most pathways identified were impacted similarly by each small molecule, with some pathways, particularly downregulated pathways, more strongly impacted by Cmp14 (**Fig. 6D, Extended Data Fig. 6E**). We identified shared downregulated genes including *SERPINE1*, *LIPG*, *NDRG1*, and *EMP1*, several of which were direct ncBAF targets (**Fig. 6E-F, Extended Data Fig. 6F-E**).

**Figure 6.**
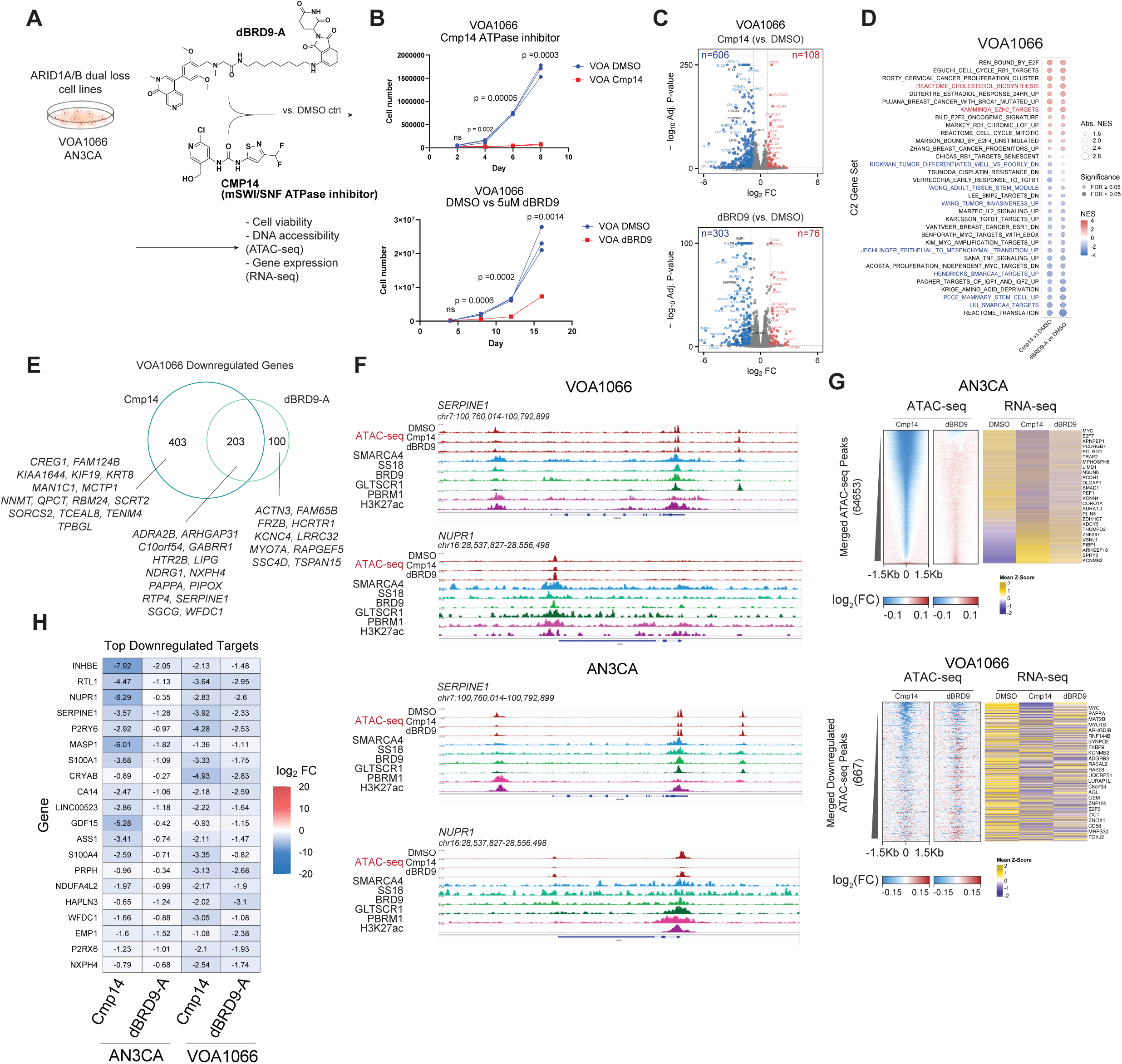
Small molecule-mediated disruption of mSWI/SNF complexes attenuates oncogenic chromatin regulation and cell proliferation. A. Schematic for small molecule inhibition experiments in DDEC cell lines in culture. B. Cell proliferation measurements for VOA1066 cells upon treatment with either Cmp14 (500nM) or dBRD9-A (1uM), compared to DMSO control. n=3 experimental replicates; Significance values from 2 sided t-test are indicated. C. Volcano plots indicating up- (red) and down- (blue) regulated genes upon either Cmp14 or dBRD9-A treatment in VOA1066 cells; 72 hours treatment, adjusted P-values calculated from DESeq2. D. GSEA performed on RNA-seq profiles in VOA1066 cells treated with Cmp14 and dBRD9-A, relative to DMSO, Adjusted P-values calculated from applying the Benjamini-Hochberg correction to GSEA P-values for all C2 gene sets. E. Venn diagram indicating overlap of downregulated genes (logFC>1; p<0.01) in Cmp14 and dBRD9-A conditions in VOA1066 cells. F. Representative ChIP-seq and ATAC-seq tracks at the SERPINE1 and NUPR1 locus in VOA1066 and AN3CA cells. G. (left) Heatmap indicating change in ATAC-seq signals in Cmp14 and dBRD9-A conditions relative to DMSO control in AN3CA and VOA 1066 cells; (right) gene expression change (z-score) from RNA-seq at selected corresponding loci shown for both cell lines. H. Top downregulated gene targets in AN3CA and VOA cell lines following Cmp14 or dBRD9-A treatment (relative to DMSO). Log2FC values are shown.

At the accessibility level, major changes were observed with the Cmp14 treatments in both cell lines whereas dBRD9-A treatment resulted in fewer accessibility changes (**Fig. 6G, Extended Data Fig. 6I-E**). Of note, Cmp14 targets the ATPase activity of both ncBAF and PBAF complexes, and in concordance, resulted in both gains and losses in accessibility (**Extended Data Fig. 6I**,L). Specifically, over sites with the most marked reduction in accessibility with Cmp14, accessibility was only modestly impacted with dBRD9-A, yet resulted in similar decreases in target gene expression (**Fig. 6G**). Examples of genes with concordant changes in accessibility and gene expression upon Cmp14 and to more modest extents, dBRD9-A, (i.e. those with decreased accessibility and decreased expression) included *RTL1*, *SERPINE1*, *NUPR1*, *EMP1*, and *S100A4* among others (**Fig. 6H, 6F**) and corresponded to EMT and stemness signatures (**Extended Data Fig. 6N**). Indeed, treatment of the ARID1A/B-dual deficient HEC1B cells resulted in greater antiproliferative impact following treatment with FHD-286, a clinical grade inhibitor of SMARCA4/2 ATPase activity^36–38^ (**Extended Data Fig. 6O-E**).

### mSWI/SNF inhibition reduces tumor growth and potentiates survival

Lastly, we established subcutaneous tumor models of the VOA1066 cell line (cell line xenograft (CDX)) and a novel PDX, XVOA14590, derived from a human DDEC primary tumor lacking both ARID1A and ARID1B expression (**Extended Data Fig. 7A**). Specifically, we evaluated either Cmp14 SMARCA4/2 ATPase inhibitor (25mg/kg dosed QD as IP injections) or the BRD9 degrader, dBRD9-A (50mg/kg, dosed QD as IP injections) (**Fig. 7A**). Treatment of XVOA14590 PDX with either Cmp14 or dBRD9-A resulted in statistically significant reductions in tumor volume by the study endpoint of 16 days, with average tumor volume for the vehicle control reaching the endpoint of 1500m^3^, while volumes for Cmp14 and dBRD9-A treatment conditions averaged 800mm^3^ and 450mm^3^ respectively (**Fig. 7B, Extended Data Fig. 7B-D**). Tumor burden was also significantly attenuated by both dBRD9-A and Cmp14 treatment (**Fig. 7C**). Degradation of BRD9 was confirmed in vivo at the study endpoint (**Extended Data Fig. 7D**). In addition, treatment of VOA1066 CDX with Cmp14 (20 mg/kg, QD, oral) significantly decreased tumor volume and final tumor weight (**Fig. 7B-C, Extended Data Fig. 7H**).

**Figure 7.**
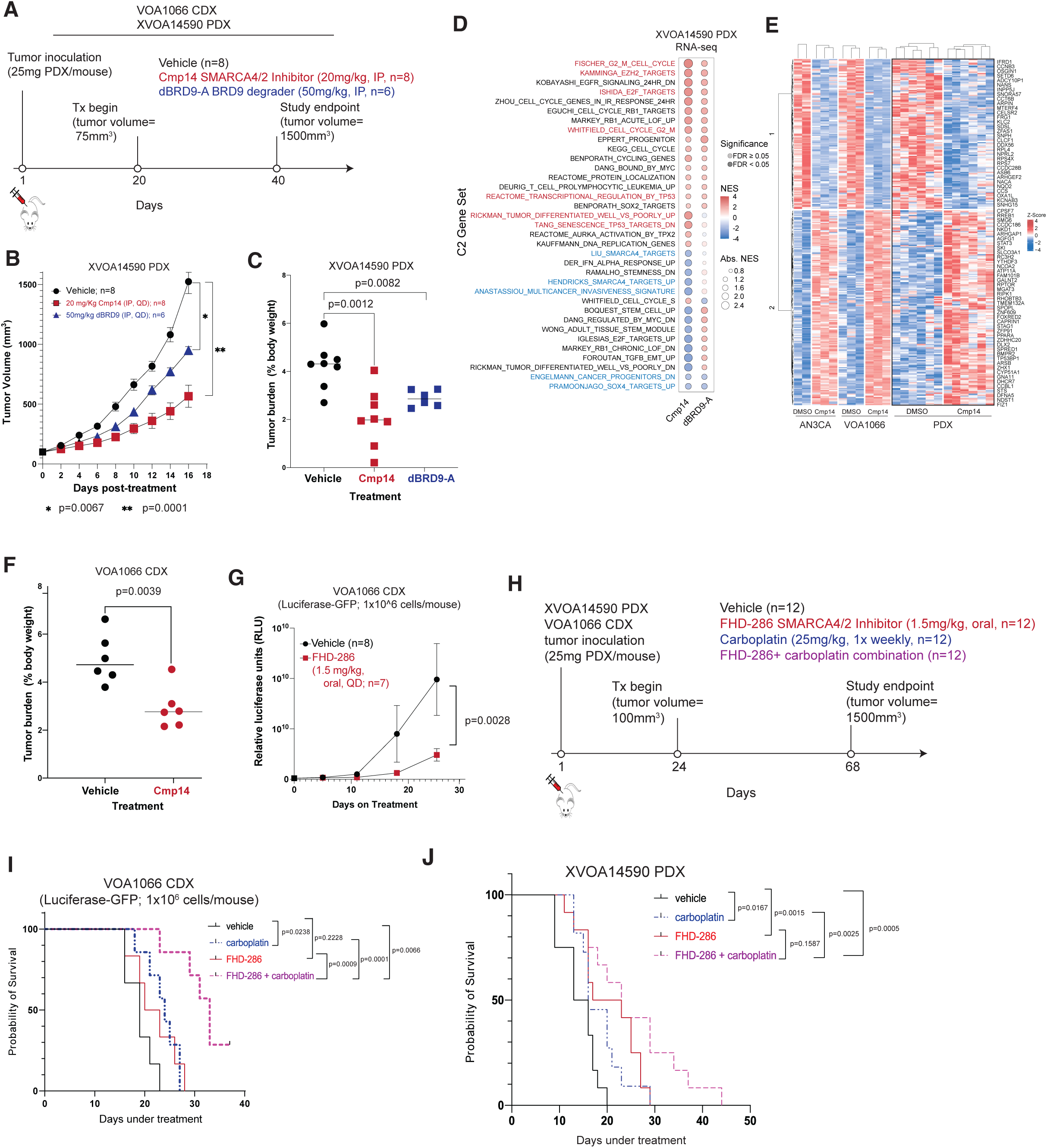
Pharmacologic disruption of residual mSWI/SNF complex activities attenuates DDEC tumor growth in vivo. A. Schematic depicting DDEC CDX and PDX tumor establishment and treatment with vehicle, Cmp14, or dBRD9-A in NRG mice. B. Tumor volume (mm3) of XVOA14590 PDX model treated with either vehicle control, 20mg/kg IP Cmp14 (QD), or 50mg/kg IP dBRD9-A (QD), p-values determined by two way ANOVA. C. Tumor burden measurements for XVOA14590 PDX model treated with either vehicle control, Cmp14, or dBRD9-A. D. GSEA C2 pathway analysis performed on RNA-seq profiles (merged; n=5) in each treatment condition relative to vehicle control in XVOA14590 PDX model at day 16 (study endpoint). NES, normalized enrichment score. Selected pathways upregulated (red) and downregulated (blue) are highlighted. E. Z-scored heatmap indicating concordantly upregulated (red) and downregulated (blue) genes in two DDEC cell lines, VOA1066 and AN3CA, and in PDX model XVOA14590 in vivo. Selected genes are labeled. F. Tumor burden measurements for VOA1066 CDX model treated with either vehicle or Cmp14. G. Relative luciferase units for VOA1066 luciferase-GFP expressing CDX model treated with either vehicle or FHD-286 over 25-day time course. H. Schematic representing CDX and PDX tumor establishment and treatment with vehicle, FHD-286 (1.5mg.kg, oral, QD), carboplatin (1.5mg/kg, IPl, weekly) and combination (FHD-286 and carboplatin) in NRG mice. I. Kaplan-Meier curves performed on VOA1066 cell line CDX model (n=12 per condition) treated with either vehicle or FHD-286. J. Kaplan-Meier curve demonstrating survival of n=12 per condition treated with conditions indicated. p-values determined with two-tailed Students t-test for C, F and log rank test for G, I and J. Error bars represent mean +/− S.E.M.

Notably, at the gene regulatory level, treatment of either Cmp14 or dBRD9-A, particularly Cmp14, led to significant suppression of several key pathways, including cancer progenitor signatures, stemness signatures, TGFB targets, SOX4 targets as well as early estrogen response, EMT and inflammatory response pathways in XVOA14590 PDX tumors (n=6) (**Fig. 7D, Extended Data Fig. 7E-G**). Pathways upregulated and downregulated only in the setting of Cmp14 treatment and not dBRD9-A included those of differentiation and restoration of P53/senescence targets (upregulated), SMARCA4 targets (downregulated) and multicancer invasiveness (downregulated), in line with the greater anti-tumor impact observed with ATPase inhibition relative to dBRD9-A in vivo (**Fig. 7D, Extended Data Fig. 7F-E**). Of note, in comparing these results with those in DDEC cell lines, we identified a collection of genes that were concordantly up- and down-regulated in 2D culture in cell line models and in PDX XVOA14590 in vivo (**Fig. 7E**). Further, Cmp14 treatment exhibited a similar anti-tumoral effect in the VOA1066 CDX model in vivo (**Fig. 7F, Extended Data Fig. 7H**).

In parallel, we evaluated the impact of FHD-286, a clinical-grade SMARCA4/SMARCA2 ATPase inhibitor ^38^. Notably, FHD-286 treatment (1.5mg/kg, QD) for 25 days also attenuated growth of VOA1066 CDX modified with GFP-luciferase (**Fig. 7G, Extended Data Fig. 7H**). Further, we sought to define whether treatment with FHD-286 would potentiate the activity of carboplatin on DDEC cell line growth in culture and tumor growth in vivo. Indeed, both ARID1A/ARID1B dual-deficient (i.e. VOA1066) as well as ARID1A-deficient models (i.e. RL95-2, SNGM) cell lines exhibited synergistic responses with combined FHD-286 and carboplatin (**Extended Data Fig. 7I**). In vivo, using the XVOA14590 PDX and VOA1066 CDX model systems, we observed significant potentiation in overall mouse survival relative to either single agent alone, absent substantial changes in body weight (**Fig. 7H-J, Extended Data Fig. 7J-E**). Together, these findings highlight the potential clinical utility of FHD-286 alone or in combination with carboplatin in DDEC.

## Discussion

Dedifferentiated endometrial carcinoma is characterized by the frequent mSWI/SNF disruption and remains a highly aggressive tumor type with poor overall survival and limited treatment options. First-line, standard-of-care treatment includes cytotoxic chemotherapy comprising carboplatin and paclitaxel and increasingly, immunotherapy, albeit with limited success and few biomarkers capable of identifying patients likely to benefit^22,39,40^. New agents that, either alone or in combination, attenuate DDEC are needed. Here we find that without these two subunits needed for cBAF assembly^15^, only residual ncBAF and PBAF mSWI/SNF remodeler subcomplexes exist on chromatin, enhancing their function via increased stoichiometric abundance and oncogenic activities. These enhanced functions uphold the cancer state, including loss of lineage-associated proteins (including cytokeratins) that are normally maintained by cBAF.

While ARID1A is widely considered a tumor suppressor in the endometrium and more broadly^41–44^, we show here that ARID1 re-expression in tumors with concomitant ARID1A/ARID1B loss promotes cBAF reassembly and binding at distal enhancers associated with epithelial differentiation and estrogen signaling pathways. Further, here, we directly restoration of ARID1A, ARID1B and both (ARID1A+ARID1B), gaining new insights into how these two paralogs shape chromatin accessibility and transcriptional landscapes (**Figure 4**). We find substantial functional redundancy between ARID1A/B paralogs but also identify gene sets relevant to oncogenic maintenance in DDEC that can only be rescued by ARID1B re-expression or the combination of ARID1A/B, opening new opportunities to define mechanisms of mSWI/SNF paralogs. Broadly, cBAF restoration leads to altered cell morphology and cytokeratin expression, phenocopying well-differentiated endometrial carcinoma and suggesting cBAF as a pivotal regulator of epithelial lineage commitment.

While the role for cBAF in regulating chromatin accessibility, particularly at distal enhancers, is well established,^13,14,18,45^ the specific functions of ncBAF and PBAF are less well understood. We previously demonstrated increased ncBAF abundance and activity in other cBAF-mutated cancer contexts including SMARCB1- and SMARCE1-deficient cancers.^18,45^ Here, our work demonstrates the intriguing persistence of both ncBAF and PBAF function in the absence of cBAF, a therapeutic vulnerability that we validate both genetically and pharmacologically. In DDEC, FHD-286 treatment proved more effective than solely targeting ncBAF with BRD9 degradation in vivo, suggesting either that the ATPase activity of both ncBAF and PBAF contribute to oncogenic maintenance, or, that BRD9 degradation is insufficient to attenuate ncBAF-mediated functions in this context. Early-phase clinical trials with FHD-286 have demonstrated tolerability with relatively favorable side effect profiles not overlapping with those of carboplatin.

In contrast to other studies,^46^ we do not find that PBAF complex functions are reduced or assembly is altered in ARID1A/B dual-deleted cancers; indeed, based on this study and prior work^15^, the cBAF assembly pathway is separate and distinct from that of PBAF and ncBAF, connected only by the fact that core modules that can no longer nucleate cBAF complexes in the absence of ARID1A/B and instead contribute to ncBAF/PBAF assembly. Of note, dual ARID1A/B loss also occurs in 4% of gastric and 1-2% of liver cancers^47–50^, further broadening the relevance of our findings. In the liver, dual ARID1A/B loss leads to similar aggressiveness and enhanced dedifferentiation coupled with impaired liver function^46^; based on our mechanistic findings, FHD-286 alone or in combination with chemotherapy may thus warrant clinical investigation in these cancers.

Until recently, mechanistic studies of endometrial carcinoma have been hindered by the scarcity of well characterized cellular models^51^. Here, the *ARID1A*/*ARID1B*-mutant cellular models used are instrumental, yet, they also have important limitations including mutations that need to be evaluated at the protein level to confirm impact (i.e. truncations in VOA1066) or DDEC histology in AN3CA (i.e. **Extended Data Fig. 1F**)^15,52^. The findings in this study are supported by datasets emerging from two distinct cell lines with differing mutational patterns with their commonality being ARID1A/ARID1B loss-of-function mutations.

In summary, this work underscores the importance of incorporating genetic profiling (e.g. ARID1A/B mutational status) in the evaluation of endometrial carcinomas and during DDEC classification. Defining DDEC by histologic subtype alone currently excludes patients from most gynecologic oncology clinical trials, including those incorporating mSWI/SNF-mutated tumors. Our data argue that mSWI/SNF-mutant DDEC/UEC lies within a spectrum of cBAF-mutated endometrial carcinomas that can be characterized by the relative abundance of remaining functional cBAF complexes, which constitutes the mechanisms and targetable dependencies defined herein. Further studies will be needed to define the consequences of specific ‘doses’ of remaining cBAF abundance in *ARID1A*-mutant endometrial carcinomas.

## Supporting information

ExtendedDataFigures

## Acknowledgements

We thank all of the members of the Kadoch Laboratory and Wang/Huntsman Laboratories for assistance and discussion. Specifically, we thank Dr. Siddhant Jain for assistance with ARID1A piggybac construct cloning, Eunice Li for isogenic cell line generation, as well as Chae Young Shin and Lien Hoang for assistance with DDEC microdissection. We also thank Zach Herbert, Maura Sullivan and Rachel Garuti of the DFCI Molecular Biology Core Facility (MBCF) for help with high-throughput sequencing. J.S.L. is supported by the NIH RDSP (Grant #K12-HD000849-37) and the GOG foundation. This work was supported in part by an NIH DP2 New Innovator Award (C.K.) and the Howard Hughes Medical Institute (HHMI), Canadian Institute of Health Research (CIHR) grant PJT-462168 and PJT-197923 (Y.W.), Terry Fox New Frontiers Program Project Grants (#1116, Y.W.; #1082, D.G.H). Y.W. is a recipient of the North Family Health Research Award. D.G.H. is supported by a Canada Research Chair in Molecular and Genomic Pathology. We also appreciate the generous support from the British Columbia Cancer Foundation and the VGH/UBC Hospital Foundation.

## Author Contributions

J.S.L: Conceptualization, methodology, validation, formal analysis, investigation, data curation, writing, visualization, project administration. G.X., A.P., D.S.G.., S.C., W.W.F., D.S.N, B.G.: Methodology, validation, formal analysis, data curation. A.W.Y., K.S.C., A.S.: Formal analysis, data curation, visualization, methodology. J.A.P.: Methodology, validation, formal analysis, software, data curation. J.Q., S.P.G: Resources. J.L.H: data curation, resources. D.J.K.: methodology, investigation, analysis, data curation, resources, manuscript editing. Y.W.: Methodology, validation, formal analysis, investigation, data curation, supervision, conceptualization-cell line screening efforts, manuscript editing, project administration, funding acquisition. D.G.H: resources, supervision, funding acquisition. C.K.: Conceptualization, methodology, investigation, resources, data curation, writing, visualization, supervision, project administration, funding acquisition.

## Methods

All research performed in this study complies with all relevant ethical regulations and was approved by institutional review boards at Dana-Farber Cancer Institute and Brigham and Women’s Hospital (BWH) (IRB 2023P003629, 2022P001059), and University of British Columbia IRB# H18-01652.

### Cell Lines

All female human cell lines (AN3CA, VOA 1066, KLE, HEC-265, HEC1A RL95-2, SNGM, HEC-50B and MFE-296 were grown at 37 °C in a humidified incubator with 5% CO_2._ KLE and RL95-2 cell lines were cultured in DMEM:F12 (Gibco) supplemented with 10% Fetal Bovine Serum (FBS) (Gibco) with 0.005mg/mL insulin (Sigma-Aldrich) for RL95-2 only. HEC1A cells were grown in McCoy’s 5A media (Gibco) with 10% FBS. HEC-265 and HEC-50B cells were grown in EMEM (Gibco) with 15% tetracycline-free FBS (Omega). SNGM cells were maintained in Ham’s F12 (Gibco) with 20% FBS. Finally, VOA1066, AN3CA, and MFE-296 were cultured in EMEM with 15% tetracycline-free FBS. HEC1B were cultured in RPMI 1640 supplemented with 5% FBS. All cell lines were maintained in 100U/mL Penicillin-Streptomycin (Gibco).

### Animal Models

All animal experiments adhered to the Canadian Council on Animal Care (CCAC) and Animals for Research Act (R.S.O. 1990, Chapter c. A.22) guidelines, with protocols approved by the University of British Columbia’s Animal Care Committee (A17-0146, A22-005). Female NRG (NOD.Rag1KO.IL2RγcKO, Jackson Labs) mice (6-8 weeks) were injected subcutaneously with either VOA1066 or VOA1066-Luciferase (luc) cells (1×10⁶ cells/mouse) or PDX tumor cells (XVOA14590, 25mg/mouse), all in a 1:1 Matrigel (Corning) mixture (final volume 200 μl). Details regarding housing conditions and drug treatments are described in the **Supplementary Note**.

### Human tumor samples

Archival paraffin embedded tissue from patients with a diagnosis of dedifferentiated, undifferentiated or well differentiated endometrial cancer were obtained under institutional IRBs: 2022P001059 (DCFI) and H18-01652 (UBC). Archived samples did not require patient consent for analysis. Diagnosis of DDEC/UDEC was established based on pathologic review by Brigham and Women’s and Vancouver General Hospital’s gynecologic pathology divisions. All cases with DNA sequencing data were previously sequenced with data obtained under research IRB 2023P003629. Deidentified variables related to clinical outcomes extracted included sex, age, stage, treatment, progression free survival and overall survival are summarized in **Supplementary Table 1**.

### Plasmids, cloning and exogenous expression

For inducible expression, ARID1A and ARID1B coding sequences were cloned downstream of a doxycycline-inducible promoter in a piggybac vector. This vector also contains a separate Tet-On 3G cassette and a Blasticidin resistance gene separated by a P2A cleavage sequence, all driven by the human EF1a promoter. Sanger sequencing verified the accuracy of all constructs. Piggybac plasmids were co-transfected with a mammalian transposase expression plasmid into AN3CA cells using TransIT-T1 Transfection Reagent (Mirus Bio Cat# MIR2300). Selection with 10 ug/ml Blasticidin for 5 days post-transfection enriched for stably transfected clones. Doxycycline treatment at 100 ng/ml for 72 hours induced expression of the transgene. A complete list of plasmids used in this study can be found in **Supplementary Table 7**.

Isogenic AN3CA and VOA1066 cancer cell lines harboring knockouts (KO) of BRD9, GLTSCR1, GLTSCR1L, SMARCA4, and SMARCA2 were generated using a commercially available CRISPR/Cas9 KO plasmid system (Santa Cruz Biotechnology). To generate inducible expression of short-hairpin RNA (shRNA) constructs targeting BRD9, ARID2 or a scrambled, non-silencing sequence were cloned downstream of a doxycycline-inducible promoter in a piggybac vector. Additional details are described in the **Supplementary Note**.

### Nuclear Extraction

Cells were harvested by trypsinization, followed by centrifugation at 125 x g for 5 minutes at 4°C. The resulting pellet was resuspended in 5 volumes of hypotonic buffer containing 50mM Tris pH 7.5, 0.1% NP-40, 1mM EDTA, 1mM MgCl_2_, 100mM NaCl, supplemented with protease inhibitors (PI) and phenylmethylsulfonyl fluoride (PMSF). After centrifugation at 1000 x g for 5 minutes at 4°C, the supernatant with cytosol was aspirated and the nuclear pellet was resuspended in EB300 buffer (50mM Tris pH 7.5, 0.1% NP-40, 1mM EDTA, 1mM MgCl_2,_ 300mM NaCl) and then incubated on ice followed by centrifugation at 21, 130 x g for 10 minutes at 4°C. Supernatant was recovered and kept at −80°C until use.

### Western Blotting

Whole cell lysates were generated using RIPA lysis buffer (50mmol/L Tris, pH 8.0, 150mmol/L NaCl, 5mmol/L MgCl2, 1% Triton X-100, 0.5% sodium deoxycholate, 0.1% SDS) supplemented with protease and phosphatase inhibitor cocktail ( Cat# 78440, Thermo Fisher). Nuclear extracts or whole cell lysates were resolved by sodium dodecyl sulfate-polyacrylamide gel electrophoresis (SDS-PAGE) using a precast 4%–12% Bis-Tris PAGE gel (Bolt 4%–12% Bis-Tris Protein Gel, Thermo Fisher Scientific) following the manufacturer’s protocol. Proteins were transferred to polyvinylidene difluoride (PVDF) membranes (0.2 μm, Bio-Rad) at 400 mA for 2 hours at 4°C. Membranes were blocked with 5% (w/v) nonfat dry milk in Tris-buffered saline with 0.1% Tween-20 (TBST) for 30 min at room temperature (RT). Membranes were then incubated with primary antibodies diluted in TBST with shaking overnight at 4°C. The primary antibody concentration used was 1:1000 (v/v) for all antibodies. After incubation, membranes were washed 3x with TBST, then probed with species-specific fluorophore-conjugated secondary antibodies (LI-COR Biosciences) at a 1:10,000 (v/v) dilution in TBST for 1 hour at RT with shaking. Membranes were washed twice with TBST followed by a final wash with Tris-buffered saline (TBS). Finally, protein bands were visualized and quantified using a Li-Cor Odyssey CLx imaging system (LI-COR Biosciences). Antibody information is contained in **Supplementary Table 8**.

### Density Sedimentation

Glycerol gradient sedimentation was performed as previously described^15^. Briefly, nuclear extracts were resuspended in EB300 buffer (50mM Tris pH 7.5, 1.0% NP-40, 1mM EDTA, 1 mM MgCl_2_, 300m NaCl) supplemented with 1 mM DTT, protease inhibitors, and 1 mM PMSF, followed by quantification using BCA. 1mg of total nuclear lysate was loaded on top of a linear 10%–30% glycerol gradients with BC0 as the base buffer. Glycerol gradient tubes were loaded into a SW41 rotor and centrifuged at 288,000 x g for 16 hours at 4°C. Following centrifugation, sample was aliquoted in 550 μL fractions of which 100-250 μL was concentrated using 10-15 μL Strataclean beads (Agilent Cat#400714) with rotation for 2 h at 4 °C. The protein-bound beads were then eluted in fresh sample buffer (2×NuPAGE LDS buffer with 100 mM DTT) and loaded onto 4–12% Bis-Tris NuPAGE Gels (Life Technologies).

### Tissue Microarray Construction and Immunohistochemistry

DDEC patient tumors were obtained from Vancouver General Hospital (VGH) with the study approval by the Institutional Review Board (IRB) at the University of British Columbia IRB# H18-01652 and from Brigham and Women’s Hospital (BWH) IRB# P001059. Informed and written consent was obtained or waived from all participants in accordance with the relevant IRB approvals. For the VGH samples following histologic review of hematoxylin and eosin (H&E) slides previously prepared for clinical use by gynecologic pathologists, duplicate 0.6 mm cores of undifferentiated and well differentiated components from formalin-fixed, paraffin-embedded tumor tissue of each case were used for tissue microarray (TMA) construction. Each TMA section was cut at 4 μm thickness onto Superfrost Plus glass slides and were processed using the Leica BOND RX Stainer. For all cases from VGH and BWH, immunohistochemical staining was performed on 4 μm formalin-fixed paraffin embedded sections with antibodies to 1:3000 ARID1A (Cat# ab182560; RRID: AB_2889973) or 1:500 ARID1A (Cat# H00057492-M01J; RRID: AB_605947), 1:250 ARID1B (Cat# H00057492-M01J; RRID: AB_605947), 1:2000 SMARCA4 (Cat# ab108318; RRID: AB_10889900), 1:200 SMARCA2 (Cat# HPA029981; RRID:AB_10602406), 1:1000 SMARCB1 (Cat# 612110; RRID: AB_399481) and 1:500 GLTSCR1 (Cat# 45441; RRID: AB_3095741). IHC staining was reviewed and scored by DK. ARID1A, ARID1B, SMARCA4, SMARCA2, and SMARCB1 were interpreted as either intact/normal if staining was present in tumor nuclei or lost/aberrant if tumor nuclei showed an absence of staining. For dedifferentiated carcinomas, ARID1A, ARID1B, SMARCA4, SMARCA2, SMARCB1 were assessed in the undifferentiated component. Appropriate internal controls showed positive staining. GLTSCR1 staining was semi-quantitatively assessed using the method described previously.^53^

### Immunofluorescence staining

VOA1066 and AN3CA cells were seeded onto sterile glass coverslips at the bottom of 6-well plates (Corning) at a density that achieved 75% confluency 72 hours after plating. To induce ARID1A expression, doxycycline (Fisher Scientific) was added to the culture medium at a final concentration of 100 ng/mL. After 72 hours, cells were washed twice with PBS followed by fixation with 3% paraformaldehyde for 20 minutes at RT. Cells were then permeabilized and blocked by incubation with Wash buffer (PBS containing 0.1% NP-40, 1 mM sodium azide, and 10% FBS) for 1 hour at RT. Pan Cytokeratin (AE1/AE3) Alexa Fluor 488 (Cat# 53-9003-82; RRID: AB_1834350) was diluted 1:200 in Wash buffer and incubated overnight at 4°C. Following incubation, cells were washed five times with Wash buffer. Finally, coverslips were mounted onto glass slides using InvitrogenProLong Gold Antifade Mountant (Fisher Scientific).

### ATAC-Sequencing

DNA accessibility was assessed using a modified Omni-ATAC protocol.^54^ 100,000 cells per sample were trypsinized and washed with cold PBS. Cell pellets were then lysed in 50 μL cold resuspension buffer (RSB) supplemented with fresh NP40 (final 0.1% v/v), Tween-20 (final 0.1% v/v), and Digitonin (final 0.01% v/v) (RSB recipe: 10 mM Tris-HCl pH 7.4, 10 mM NaCl, and 3 mM MgCl2). This lysis step was quenched with 1 mL of RSB supplemented with Tween-20 (final 0.1% v/v), and nuclei were pelleted at 500 x g for 10 min at 4 °C after incubating on ice for 3 minutes. Nuclei were then resuspended in 50 μL transposition reaction mix containing 25 μL 2X Tagment DNA buffer (Illumina Cat#20034198), 2.5 μL Tn5 transposase (Illumina Cat#81286), 16.5 μL 1X PBS, 0.5 μL 1% digitonin (final 0.01% v/v), 0.5 μL 10% Tween-20 (final 0.1% v/v), and 5 μL of nuclease-free water. The transposition reaction was carried out at 37 °C for 30 min with constant shaking (100 x g) on a thermomixer. Tagmented DNA was purified using the MinElute Reaction Cleanup Kit (Qiagen Cat#28204). 7 cycles of amplification was used to amplify tagmented libraries.^55^ Libraries were sequenced on a NextSeq 500 (Illumina) using 37 bp paired-end sequencing.

### RNA-Sequencing

For bulk cell line RNA sequencing, total RNA was isolated from two million cells per sample of AN3CA and VOA1066 cell lines; all experimental conditions were performed in triplicate. Cells were first collected and washed with cold PBS to remove trypsin, then stored in RLT buffer (Qiagen) for subsequent RNA isolation. RNA purification was performed using the RNeasy kit (Qiagen Cat#74106) following the manufacturer’s instructions. Next, RNA libraries were prepared using the NEBNext Ultra II Directional RNA Library Prep Kit (Illumina Cat#E7760L). The quality and quantity of these libraries were assessed using a TapeStation (Agilent) and a Qubit Fluorometer (Thermo Fisher Scientific), respectively. Finally, the prepared RNA-seq libraries were sequenced on an Illumina NextSeq 500 platform using a single-end, 75-bp read length. For RNA extracted from FFPE tissue, please see **Supplementary Note**.

### ChIP-Sequencing

Chromatin Immunoprecipitation (ChIP) was performed on 40 million fixed cells. Cells were trypsinized, washed twice with PBS, and then divided into aliquots and crosslinked with 1% formaldehyde for 10 minutes at 37°C with constant agitation. Crosslinking was quenched with glycine, and cells were then washed with cold PBS before storage at −80°C in 10 million cell aliquots. Ten million cells were used per epitope for subsequent ChIPs. Following nuclei isolation, chromatin was sheared to a desired size range using a Covaris E220 sonicator. Sonicated chromatin was clarified by centrifugation, and the supernatant was incubated overnight at 4°C with specific antibodies (2–3 μg) (see antibody list). Protein G Dynabeads (Thermo Fischer Cat#10004D) were used to capture antibody-chromatin complexes, which were then washed extensively. Following elution from beads, eluate was treated with RNase A and Proteinase K to remove RNA and proteins, respectively. Purified ChIP DNA was isolated using SPRI beads, washed, and eluted in low-TE buffer for storage at −20°C. ChIP DNA libraries were prepared using the Illumina NEBNext Ultra II DNA Library Prep Kit following the manufacturer’s instructions. Finally, all libraries were sequenced using single-end 75bp reads on the Illumina NextSeq 500 platform.

### Laser Capture Microdissection and RNA sequencing from primary tumor specimens

For the isolation of well and dedifferentiated cancer components, laser capture microdissection (LCM) was employed. Briefly, tissue sections (5 μm) were cut from fixed paraffin embedded tissue blocks and mounted on PEN membrane Frame Slides (Thermo Fischer Cat# LCM0521). Paraffin-embedded sections were deparaffinized and rehydrated before H&E staining as follows. Briefly, slides were fixed in xylene for two minutes, graded ethanol from 100-75% for a total of three minutes, water for 2 minutes, hematoxylin for 2 minutes, water for 2 minutes, graded ethanol for 3 minutes and xylene for 1 minute then air-dried. Under a microscope integrated with the LCM system (Leica Microsystems), tumor cells were identified based on regions selected on hematoxylin and eosin slides with a goal of 5,000 cells per sample. Subsequently, a laser beam was used to precisely dissect the desired regions of interest, capturing the isolated cancer cells onto the CapSure film (Thermo Fischer Cat#A30155) subjected to RNA isolation.

### Small molecule treatments for cell proliferation and genomics assays

To induce expression of ARID1A or ARID1B transgenes AN3CA and VOA1066 cells were grown in MEM media either supplemented with 200ng/mL Doxycycline to induce expression for 72 hours before harvesting for ATAC-seq, ChIP-seq and RNA-seq. For drug treatment studies, AN3CA and VOA1066 cells were treated with DMSO, 500-750nM Compound 14 or 1µM dBRD9a for 72 hours before harvesting for ATAC-seq, ChIP-seq and RNA-seq (see Methods below).

### Cell growth and viability assays

Cell viability assays and dose-response curves were performed for VOA1066, RL-952, SNGM, and AN3CA cell lines in 96-well plates with three technical replicates per condition. For proliferation assays, cells were seeded at 500 cells per well and treated for 6 days with drugs suspended in DMSO. The drug-containing media was replaced every 3 days with fresh media containing the corresponding drug concentration. Following the treatment period, cell viability was measured using a fluorescence-based viability assay, CellTiter-Glo® (Promega Cat#G9242). Synergy between drugs was evaluated using the Combenefit software as previously described. Cells were plated in a 6×6 drug concentration matrix format in triplicate and viability was measured after 6 days using CellTiter-Glo®. Bliss synergy scores were calculated for each drug combination and mapped relative to cell proliferation using Combenefit v2.021. Colony formation assays were performed in 6-well plates. 1,000 cells per well were seeded and treated with 100 ng/mL doxycycline or indicated drugs for 14 days. After treatment, cells were washed with PBS, fixed with 4% paraformaldehyde, and stained with Crystal Violet solution (Sigma Aldrich Cat# V5265).

### HEC1B and HEC1B.KO CRISPR screen

HEC1B cells were transiently transfected with a pool of 3 GFP-tagged CRISPR/Cas9 KO plasmids targeting ARID1B (Santa Cruz, sc-402365). GFP-positive cells were sorted out for clonal expansion and screen of ARID1B knockout clones. To perform genome-wide dropout screen, Both HEC1B and an ARID1B knockout clone (HEC1B.BKO) cells were infected with TKOv3 genome-wide library (Addgene, #90294; PMID: 28655737) at a transduction efficiency of 30% aiming 500 times representation of each sgRNA. Cells were selected with puromycin for two days and 25×10^6 cells collected as a baseline. Remaining cells were kept in culture for 18 days post-infection and collected as the endpoint. Genomic DNA was isolated using QIAamp DNA Blood Maxi Kit following manufacturer’s instructions. sgRNA library was amplified from genomic DNA using Illumina-compatible index primers and sequenced by NovaSeq platform aiming for 300-500 times read coverage. The sequencing data was analyzed statistically using the MAGeCK software to identify dependencies.^56^

### Tandem Mass Tag (TMT)-Mass Spectrometry Data Analysis

Differential abundance analysis was performed on normalized protein counts using edgeR’s glmFit and glmLRT functions (Figure 3C, S3G). Obtained log fold-change values and adjusted p-values were visualized using ggplot2. Log2FC values were calculated based on normalized protein counts and were plotted.

### RNA-seq Data Analysis

RNA-Seq reads were demultiplexed using bcl2fastq v2.20.0.422 aligned to the hg19 genome with STAR v2.5.2b.^57^ Upregulated and downregulated genes were determined using DESeq2 (log2FC = 1, B-H p-value = 0.05).^58^ DESeq2’s estimateSizeFactors function was used to generate normalized counts. ggplot2^59^ was used to generate volcano plots for changes in expression visualized as scatter plots. The eulerr R package^60^ was used to generate venn diagrams of differential genes. Variance stabilizing transformed counts were used to generate PCA plots across the top 500 genes and plotted using ggplot2^61^. ComplexHeatmap R package^62^ was used to generate heatmaps and visualize Z-Score normalized read counts for each gene across all conditions for each cell line. The Kendall and Spearman distance functions were used to perform hierarchical clustering in heatmaps. clusterProfiler GSEA function^63^ was used through the msigdbr R package to perform GSEA analysis using the Hallmark and C2 gene sets. The “stat” output from DESeq2 was used to determine Differential expression ranking. For GSEA analysis in Fig S3D, GSEA software was obtained from the Gene Set Enrichment Analysis website [http://www.broad.mit.edu/gsea/downloads.jsp].

### ATAC-seq Data Analysis

ATAC-seq reads were first trimmed using Trimmomatic v0.36, aligned with Bowtie2 v2.2.9, and filtered with Picard v2.8.0 (MarkDuplicates REMOVE_DUPLICATES=true) and SAMtools v 0.1.19 (-F 256 -f 2 - q 30). Reads mapping to regions defined in the ENCODE project’s wgEncodeDacMapabilityConcensusExcludeable bed file were removed using bedtools v2.30.0. ATAC-seq data was processed by merging technical replicates using SAMtools merge. Peaks were called using MACS3 (-f BAMPE -g hs -q 0.001 --nomodel --extsize 200 –broad). Differential accessibility analyses are detailed in the **Supplementary Note**.

### ChIP-seq Data Analysis

ChIP-seq reads were aligned in a manner identical to ATAC-seq reads. Peaks were generated with MACS2 (AN3 CA: -f BAM -g hs –broad –broad-cutoff 0.01 --nomodel --extsize 150, VOA1066: -f BAM -g hs -q 0.001 --nomodel) Differential binding was determined using bedtools intersect and visualized using matplotlib. Tracks were generated using deepTools bamCoverage (--binSize “40” --normalizeUsing “CPM” --exactScaling). Additional information regarding ChIP-seq analyses are detailed in the **Supplementary Note**.

### Statistics and Reproducibility

GraphPad Prism 8 software was used to generate graphs and statistical analysis. Statistical significance was determined by Student’s t-test, one- and two-way ANOVA, and log-rank test, as appropriate. All experiments have been repeated in at least two independent experiments (n=2 biologic replicates). No data were excluded from the analyses. All immunoblots are representations of independent experiments.

## Data Availability

All sequencing raw and processed data are available in the Gene Expression Omnibus (GEO) database under the series GSE268960; http://www.ncbi.nlm.nih.gov/geo/query/acc.cgi?acc=G SE268960 or in the European Genome-phenome Archive (EGA) under study ID: EGAS50000001004; http://www.ega-archive.org/datasets/. All raw proteomics and mass-spectrometry datasets have been deposited to ProteomeXchange via PRIDE database under accession number PXD058236.

## Code availability statement

No new code was generated for this work; all analyses have been performed according to the details described in the Methods (RNA-seq Data Analysis, ATAC-seq Data Analysis, ChIP-seq Data Analysis, Tandem Mass Tag (TMT)-Mass Spectrometry Data Analysis, and Statistics and Reproducibility sections) using available software packages.

