## ExtendedDataFigures for "Shifted assembly and function of mSWI/SNF family subcomplexes underlie targetable dependencies in dedifferentiated endometrial carcinomas"

**St. Laurent et al., 2025**

**Extended Data Figures and Legends**

| Figure | Figure Title | Figure Legend |
|---|---|---|
| **Extended Data Fig. 1** | **mSWI/SNF gene mutations in dedifferentiated endometrial carcinomas.** | A. Summary of whole-exome sequencing profiling of uterine dedifferentiated carcinomas (GENIE cohort, n=22 cases), indicating mutation frequencies and type of mSWI/SNF, POLE, TP53, and other gene mutations. B. Summary of whole-exome sequencing profiling of endometrial/ovarian dedifferentiated carcinomas (Dana Farber dataset; n=35 cases), indicating mutation frequencies and type of mSWI/SNF, POLE, TP53, and other genes. C. Summary of mutations in TCGA (n=515 cases; serous/mixed/endometrioid carcinoma). Features are indicated in legend; Tumor type, tumor grade, mutation type and histology are indicated. D. Pie chart showing breakdown of mSWI/SNF gene mutations in DDEC cases from DFCI, Coatham et al., and TCGA. E. Mutual exclusivity analyses performed for mSWI/SNF, POLE, and TP53 on TCGA dataset of serous/mixed/endometrioid dedifferentiated carcinomas. F. Mutational characteristics for endometrial carcinoma cell lines used in this study. mSWI/SNF and top-mutated genes are shown (CCLE). *indicates damaging mutations. G. Immunoblot performed on nuclear extract (input) and IgG and anti-SMARCA4 immunoprecipitation from VOA1066 cells. mSWI/SNF components and TBP are shown. H. TMT-MS shown for SMARCA4 IP (and IgG control IP) in VOA1066 cells. I-J. Summary of n=75 DDEC cases analyzed by IHC, indicating positive/negative protein staining status of ARID1A, ARID1B, SMARCA4, SMARCA2, and SMARCB1. K. SMARCA4 IHC performed on n=3 ARID1A/ARID1B-dual mutated DDEC cases. Undifferentiated and well-differentiated compartments are indicated. |
| **Extended Data Fig. 2** | **cBAF reassembly controls genome-wide redistribution of mSWI/SNF distribution and chromatin accessibility in DDEC cells.** | A. Density sedimentation analyses performed on AN3CA ARID1A/B dual-deficient cells in +Dox (ARID1A rescue) conditions using 10-30% glycerol gradients. Immunoblot for selected mSWI/SNF subunits are shown; -Dox pair shown in Figure 1E. B. PCA plot showing -/+ Dox (ARID1A rescue) conditions in AN3CA and VOA1066 cells. C. Volcano plots showing proteins (peptide signal) significantly increased and decreased upon rescue |

| | | |
|---|---|---|
| | | with ARID1A in AN3CA and VOA cell line systems. mSWI/SNF complex components are highlighted. D. Clustered heatmap demonstrating logFC of TMT mass-spectrometry signal in AN3CA and VOA1066 cells by subunit. E. Immunofluorescence imaging for pan-cytokeratin (green) with DAPI counterstain in both DDEC cell lines in -/+ dox conditions; scale bar= 20um, Dox treatment of 72 hours, representative of n=3 experimental replicates. F. Colony formation assays performed in AN3CA and VOA1066 cell lines following 14 days of Dox treatment (ARID1A rescue). G. Quantification of colony formation assays shown in (F), p-values determined with two-tailed Students t-test, error bars represent mean +/- S.E.M. H. Bar graphs depicting number of peaks across antibodies profiled in ChIP-seq experiments. Venn diagrams depicting BRD9-GLTSCR1 (ncBAF) merged peaks in -/+ ARID1A rescue conditions in AN3CA and VOA1066 cell lines. I. Distance-to-TSS stacked bar graphs indicating peak localization for gained, lost, and retained SMARCA4 sites. J. HOMER motif analyses performed over gained and lost SMARCA4 sites in VOA1066 cells. K. PCA plot performed on ATAC-seq (broad peaks) datasets in -/+ Dox (ARID1A rescue) conditions in AN3CA and VOA1066 cells. L. Bar graphs depicting number of ATAC-seq peaks in AN3CA and VOA1066 cells in both conditions. M. Heatmaps reflecting ChIP-seq for GLTSCR1 and BRD9 as well as ATAC-seq in VOA1066 over ATAC-seq sites. Gained, lost and retained peak sets are indicated. N. Top, Overlap between de novo gained ARID1A peaks and BRD9/GLTSCR1 peaks prior to ARID1A restoration. Bottom, pie charts reflecting percent of ncBAF target sites replaced by cBAF following ARID1A restoration. O. Metaplot showing altered PBRM1 occupancy over TSS sites (top 20% expressed genes) in -/+ dox conditions. P. PBRM1 peaks in VOA1066 cells in -/+ dox conditions. |
| **Extended Data Fig. 3** | **Gene regulatory impact of cBAF restoration in DDEC cell lines and primary tumors.** | A. Immunoblot for indicated mSWI/SNF components following ARID1A restoration (72 hours). B. PCA plots performed on RNA-seq in AN3CA and VOA1066 cell lines following ARID1A restoration (n=3 experimental replicates). C. Heatmaps indicating clustering of -Dox and +Dox (+HA-ARID1) conditions. D. GSEA analyses performed on AN3CA and VOA cells (ARID1A rescue/no rescue); NES, Normalized Enrichment |

| | | |
|---|---|---|
| | | Score. E. Clustered heatmap indicating genes up and downregulated following ARID1A rescue. F. Venn diagrams indicating cell line-specific and shared up- and down-regulated genes upon ARID1A rescue. G. Volcano plots showing up- (red) and down- (blue) regulated proteins (by TMT-MS) following 72 hours of ARID1A restoration in AN3CA and VOA1066 DDEC cell lines. H. Representative ChIP-seq and ATAC-seq tracks at the CDH1, CLDN4, TONSL, and BRCA2 gene loci with corresponding bar graphs depicting gene expression (RNA-seq). |
| **Extended Data Fig. 4** | **Concordant and distinct impacts of ARID1A, ARID1B, or dual paralog restoration in DDEC cell lines.** | A. Immunoblot for indicated mSWI/SNF components following ARID1A, ARID1B or dual ARID1A/B restoration (72 hours). B. Bar graphs indicating quantitative densitometry of blots in (I). C. Venn diagrams indicating overlap of downregulated genes (left) and sites with decreased accessibility (right) following ARID1A, ARID1B or ARID1A/B dual rescue. D. GSEA (C2 gene set) performed on AN3CA cells with ARID1A, ARID1B or dual paralog rescue relative to no rescue (-Dox) in AN3CA cells. E. Z-score heatmap indicating genes upregulated, downregulated and corresponding genes in undifferentiated (UD) and well-differentiated (WD) primary tumor compartments. F. Hallmark GSEA in AN3CA, VOA1066 cell lines following ARID1A rescue (relative to - Dox) and DDEC UD tumor compartment, normal endometrium relative to DDEC WD tumor compartment (patient); NES, Normalized Enrichment Score. |
| **Extended Data Fig. 5** | **Cell proliferation assays and CRISPR screening in cBAF-disrupted endometrial cell lines.** | A. Colony formation assays performed on AN3CA cells bearing shRNAs targeting either BRD9 or ARID2. B. Immunoblot for ARID2, BRD9 and GAPDH in VOA1066 cells treated with indicated shRNA conditions. C. Bar graph depicting AUC values in indicated conditions in VOA1066 colony formation assays. D. Immunoblot for shRNA-mediated suppression of subunits indicated in AN3CA cells. E. Bar graph depicting AUC values in indicated conditions in AN3CA colony formation assays. F. Normalized read counts from genome-scale CRISPR screen (TKOv3 library). G. Box and whisker plots indicating impact across gene sets indicated on X-axis. H. Colony formation assays performed on HEC1B cells and HEC1B cells with ARID1B KO (ARID1B-BKO) treated with the indicated sgRNAs. I. Quantification of colony |

| | | |
|---|---|---|
| | | formation assays from (H) (relative confluency) at Day 10 post infection. J. Colony formation assays performed on HEC1B cells and HEC1B cells with ARID1B KO (ARID1B-BKO) treated with the indicated sgRNAs. K. Quantification of colony formation assays from (J) (relative confluency) at Day 10 post infection. L. Venn diagram depicting overlap between HEC1B ARID1B KO-specific dependencies and genes downregulated by ARID1 paralog restoration in AN3CA cell lines. p-values determined with two-tailed Students t-test for C, E, I and K. Error bars represent mean +/- S.E.M. |
| **Extended Data Fig. 6** | **SMARCA4/2 ATPase inhibition and BRD9 degradation attenuate oncogenic gene regulation and proliferation in DDEC cells.** | A. Cell proliferation measurements for VOA1066 cells upon treatment with either Cmp14 or dBRD9-A, compared to DMSO control. n=3 experimental replicates; Significance values are indicated. B. Principal component analysis (PCA) performed on RNA-seq in VOA1066 and AN3CA cells in conditions indicated. C. Volcano plots indicating up- (red) and down- (blue) regulated genes upon either Cmp14 or dBRD9-A in VOA1066 cells; 72 hours treatment. D. GSEA Hallmark performed on RNA-seq profiles in VOA1066 cells treated with Cmp14 and dBRD9-A, relative to DMSO. E. GSEA C2 gene set analysis performed in AN3CA cells treated with the indicated conditions. F. Venn diagram indicating overlap of upregulated genes (logFC>1; p<0.01) in Cmp14 and dBRD9-A conditions in VOA1066 cells. G. Venn diagram indicated overlap of downregulated (top) and upregulated (bottom) genes (logFC>1; p<0.01) in AN3CA cells. H. Representative ChIP-seq and ATAC-seq tracks at the GDF15 and EMP1 loci. I. PCA performed on ATAC-seq data in AN3CA and VOA1066 cells. J. Venn diagrams showing overlap of ATAC-seq peaks in AN3CA and VOA1066 cells in conditions indicated. K. HOMER motif analysis over sites with retained acccessibility (no change) in AN3CA and VOA1066 cells with dBRD9-A conditions. L. Venn diagrams indicating overlap between accessible peaks (ATAC-seq) in DMSO and Cmp14 conditions in each cell line. M. Homer motif analyses over sites with lost accessibility upon Cmp14 treatment. N. GSEA C2 gene set analysis of genes with altered accessibility withiin 2kB in AN3CA cells treated with either Cmp14 or dBRD9-A. O. Graph indicating relative confluency of HEC1B and HEC1B ARID1B KO cells across concentrations of FHD-286. P. Colony formation |

| | | |
|---|---|---|
| | | assays performed on HEC1B and HEC1B ARID1B KO cells treated with FHD-286 across concentrations indicated. |
| **Extended Data Fig. 7** | **mSWI/SNF pharmacologic inhibition attenuates DDEC tumor growth in vivo.** | A. H&E and IHC for ARID1A, ARID1B, SMARCA4 and SMARCA2 in the XVOA14590 PDX model system; scale bars 200um. B. Individual mouse tumor volume measurements (mm3) in XVOA14590 PDX model treated with either Cmp14 or dBRD9-A. C. Representative tumor images for XVOA14590 PDX model treated with vehicle control, Cmp14, or dBRD9-A; tumors isolated at Day 16, end of study. D. Immunoblot performed on whole-cell extracts for BRD9, CRBN and beta-actin in XVOA14590 PDX treated with dBRD9-A confirming degradation relative to vehicle. E. PCA analysis performed on RNA-seq profiles derived from n=5 tumors in each condition indicated. F. Volcano and dot plots representing differentially expressed genes in Cmp14 vs DMSO cell lines (left) and dBRD9-A treated cells (right). G. GSEA Hallmark analyses performed on n=6 primary tumors from XVOA14590 at Day 16 post treatment in vivo. H. Tumor weight measurements (g) for VOA1066 CDX treated with either Cmp14 (top) or FHD-286 (bottom). I. (Left) Cell proliferation experiments of VOA, RL95-2 and SNGM cells (by % confluency) using single-agent and FHD-286 and carboplatin combination. (Right) D-R-(LOWE) synergy plots using Combenefit studies performed in 2D culture. J. Percent change in body weight for XVOA14590 PDX model over days of treatment with single-agent and FHD-286 and carboplatin combinations. K. Individual mouse tumor volume measurements (mm3) in VOA1066 CDX model treated with vehicle, single-agent FHD-286, carboplatin, or combination FHD-286+carboplatin. |

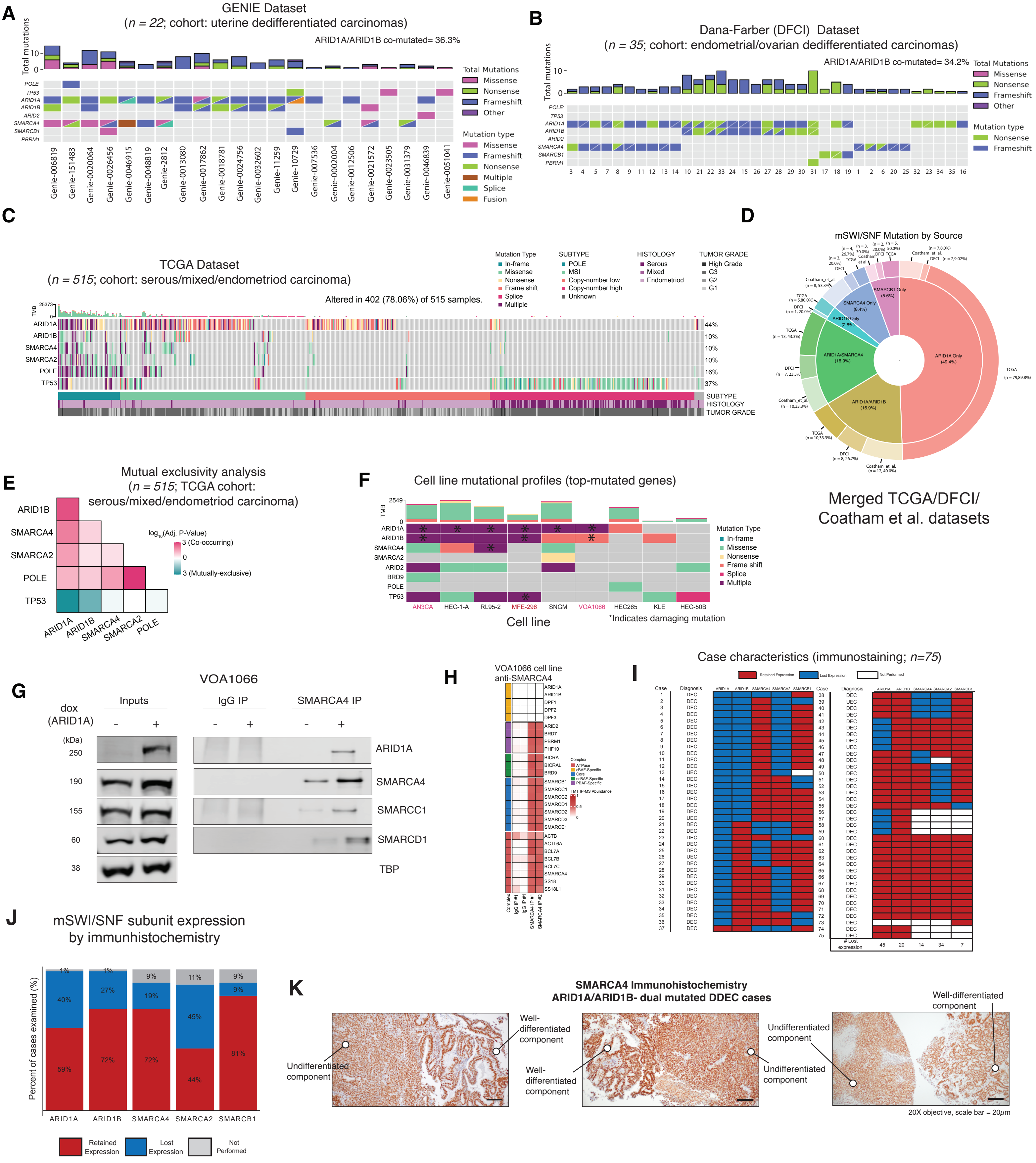
A
GENIE Dataset
(n = 22; cohort: uterine dedifferentiated carcinomas)
ARID1A/ARID1B co-mutated= 36.3%
Total mutations
POLE
TP53
ARID1A
ARID1B
ARID2
SMARCA4
SMARCB1
PBRM1
Total Mutations
Missense
Nonsense
Frameshift
Other
Mutation type
Missense
Frameshift
Nonsense
Multiple
Splice
Fusion
Genie-006819
Genie-151483
Genie-0020064
Genie-0026456
Genie-0046915
Genie-0048819
Genie-2812
Genie-0013080
Genie-0017862
Genie-0018781
Genie-0024756
Genie-0032602
Genie-11259
Genie-10729
Genie-007536
Genie-0002004
Genie-0012506
Genie-0021572
Genie-0023505
Genie-0031379
Genie-0046839
Genie-0051041
B
Dana-Farber (DFCI) Dataset
(n = 35; cohort: endometrial/ovarian dedifferentiated carcinomas)
ARID1A/ARID1B co-mutated= 34.2%
Total Mutations
Missense
Nonsense
Frameshift
Other
Mutation type
Nonsense
Frameshift
C
TCGA Dataset
(n = 515; cohort: serous/mixed/endometriod carcinoma)
Altered in 402 (78.06%) of 515 samples.
Mutation Type
In-frame
Missense
Nonsense
Frame shift
Splice
Multiple
SUBTYPE
POLE
MSI
Copy-number low
Copy-number high
Unknown
HISTOLOGY
Serous
Mixed
Endometrioid
TUMOR GRADE
High Grade
G3
G2
G1
ARID1A 44%
ARID1B 10%
SMARCA4 10%
SMARCA2 10%
POLE 16%
TP53 37%
SUBTYPE
HISTOLOGY
TUMOR GRADE
D
mSWI/SNF Mutation by Source
ARID1A Only (49.4%)
ARID1A/ARID1B (16.9%)
ARID1A/SMARCA4 (16.9%)
ARID1B Only (2.8%)
SMARCA4 Only (8.4%)
SMARCB1 Only (5.6%)
Merged TCGA/DFCI/
Coatham et al. datasets
E
Mutual exclusivity analysis
(n = 515; TCGA cohort:
serous/mixed/endometriod carcinoma)
ARID1B
SMARCA4
SMARCA2
POLE
TP53
ARID1A
log10(Adj. P-Value)
3 (Co-occurring)
0
3 (Mutually-exclusive)
F
Cell line mutational profiles (top-mutated genes)
Mutation Type
In-frame
Missense
Nonsense
Frame shift
Splice
Multiple
AN3CA
HEC-1-A
RL95-2
MFE-296
SNGM
VOA1066
HEC265
KLE
HEC-50B
Cell line
*Indicates damaging mutation
G
VOA1066
dox
(ARID1A)
Inputs
IgG IP
SMARCA4 IP
(kDa)
250
190
155
60
38
ARID1A
SMARCA4
SMARCC1
SMARCD1
TBP
H
VOA1066 cell line
anti-SMARCA4
I
Case characteristics (immunostaining; n=75)
Retained Expression
Lost Expression
Not Performed
### Lost expression
45
20
14
34
7
J
mSWI/SNF subunit expression
by immunhistochemistry
Percent of cases examined (%)
ARID1A
ARID1B
SMARCA4
SMARCA2
SMARCB1
Retained Expression
Lost Expression
Not Performed
K
SMARCA4 Immunohistochemistry
ARID1A/ARID1B- dual mutated DDEC cases
Undifferentiated component
Well-differentiated component
20X objective, scale bar = 20μm

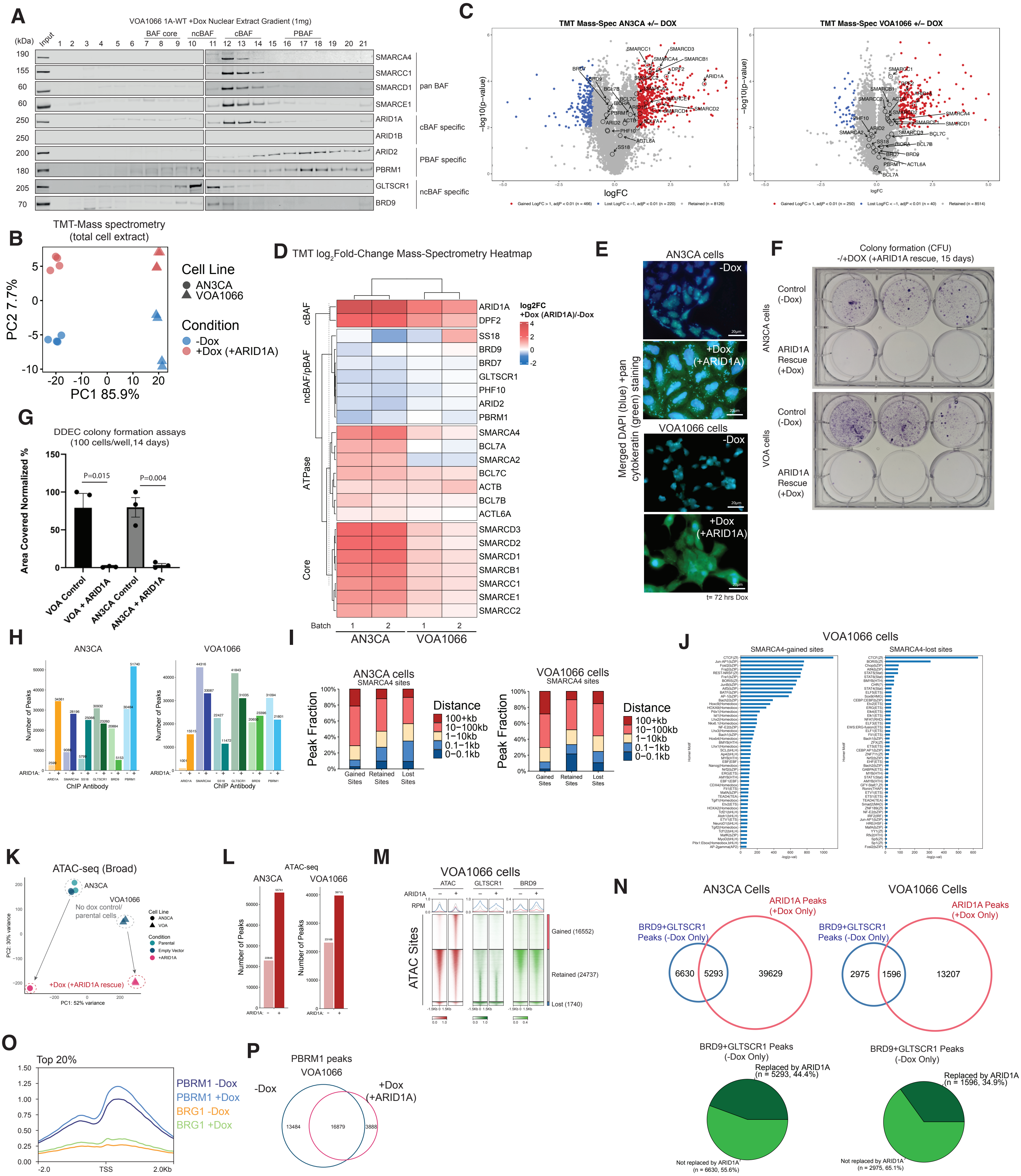
A
VOA1066 1A-WT +Dox Nuclear Extract Gradient (1mg)
(kDa)
Input
BAF core
ncBAF
cBAF
PBAF
SMARCA4
SMARCC1
SMARCD1
SMARCE1
pan BAF
ARID1A
ARID1B
cBAF specific
ARID2
PBRM1
PBAF specific
GLTSCR1
BRD9
ncBAF specific
C
TMT Mass-Spec AN3CA +/− DOX
TMT Mass-Spec VOA1066 +/− DOX
−log10(p−value)
logFC
B
TMT-Mass spectrometry
(total cell extract)
PC2 7.7%
PC1 85.9%
Cell Line
AN3CA
VOA1066
Condition
-Dox
+Dox (+ARID1A)
D
TMT log2Fold-Change Mass-Spectrometry Heatmap
log2FC
+Dox (ARID1A)/-Dox
cBAF
ncBAF/pBAF
ATPase
Core
ARID1A
DPF2
SS18
BRD9
BRD7
GLTSCR1
PHF10
ARID2
PBRM1
SMARCA4
BCL7A
SMARCA2
BCL7C
ACTB
BCL7B
ACTL6A
SMARCD3
SMARCD2
SMARCD1
SMARCB1
SMARCC1
SMARCE1
SMARCC2
Batch
AN3CA
VOA1066
E
AN3CA cells
VOA1066 cells
-Dox
+Dox
(+ARID1A)
Merged DAPI (blue) +pan cytokeratin (green) staining
t= 72 hrs Dox
F
Colony formation (CFU)
-/+DOX (+ARID1A rescue, 15 days)
Control
(-Dox)
ARID1A
Rescue
(+Dox)
AN3CA cells
VOA cells
G
DDEC colony formation assays
(100 cells/well,14 days)
Area Covered Normalized %
P=0.015
P=0.004
VOA Control
VOA + ARID1A
AN3CA Control
AN3CA + ARID1A
H
AN3CA
VOA1066
Number of Peaks
ChIP Antibody
I
AN3CA cells
SMARCA4 sites
VOA1066 cells
SMARCA4 sites
Peak Fraction
Distance
100+kb
10−100kb
1−10kb
0.1−1kb
0−0.1kb
Gained Sites
Retained Sites
Lost Sites
J
VOA1066 cells
SMARCA4-gained sites
SMARCA4-lost sites
K
ATAC-seq (Broad)
AN3CA
VOA1066
No dox control/
parental cells
+Dox (+ARID1A rescue)
L
ATAC-seq
AN3CA
VOA1066
Number of Peaks
M
VOA1066 cells
ATAC
GLTSCR1
BRD9
ATAC Sites
Gained (16552)
Retained (24737)
Lost (1740)
N
AN3CA Cells
VOA1066 Cells
ARID1A Peaks
(+Dox Only)
BRD9+GLTSCR1
Peaks (-Dox Only)
6630
5293
39629
2975
1596
13207
BRD9+GLTSCR1 Peaks
(-Dox Only)
Replaced by ARID1A
(n = 5293, 44.4%)
Not replaced by ARID1A
(n = 6630, 55.6%)
Replaced by ARID1A
(n = 1596, 34.9%)
Not replaced by ARID1A
(n = 2975, 65.1%)
O
Top 20%
PBRM1 -Dox
PBRM1 +Dox
BRG1 -Dox
BRG1 +Dox
-2.0
TSS
2.0Kb
P
PBRM1 peaks
VOA1066
-Dox
+Dox
(+ARID1A)
13484
16879
3888

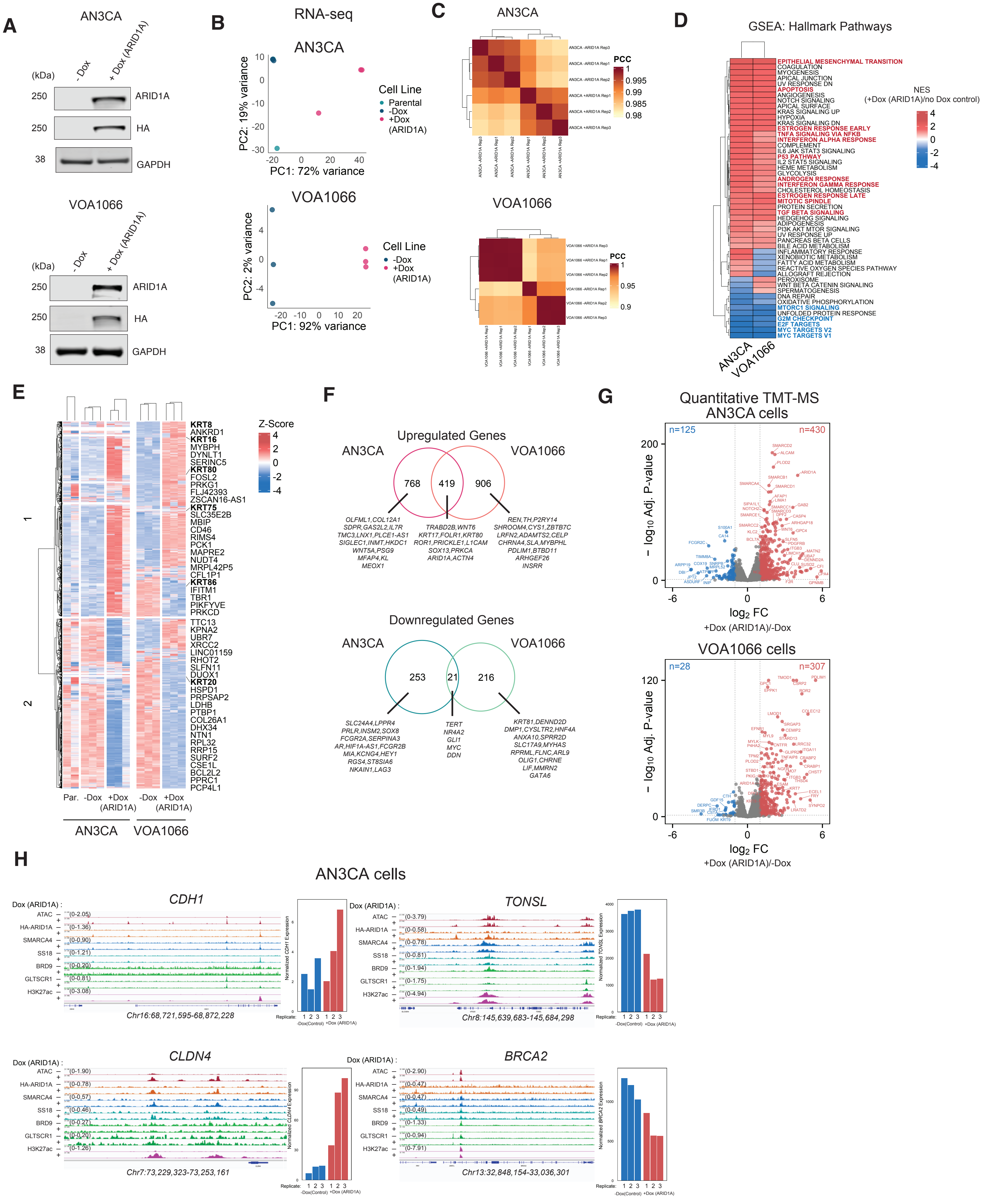
A
AN3CA
- Dox
+ Dox (ARID1A)
(kDa)
250 ARID1A
250 HA
38 GAPDH
VOA1066
B
RNA-seq
AN3CA
PC2: 19% variance
PC1: 72% variance
Cell Line
Parental
-Dox
+Dox (ARID1A)
VOA1066
PC2: 2% variance
PC1: 92% variance
C
PCC
D
GSEA: Hallmark Pathways
NES (+Dox (ARID1A)/no Dox control)
EPITHELIAL MESENCHYMAL TRANSITION
COAGULATION
MYOGENESIS
APICAL JUNCTION
UV RESPONSE DN
APOPTOSIS
ANGIOGENESIS
NOTCH SIGNALING
APICAL SURFACE
KRAS SIGNALING UP
HYPOXIA
KRAS SIGNALING DN
ESTROGEN RESPONSE EARLY
TNFA SIGNALING VIA NFKB
INTERFERON ALPHA RESPONSE
COMPLEMENT
IL6 JAK STAT3 SIGNALING
P53 PATHWAY
IL2 STAT5 SIGNALING
HEME METABOLISM
GLYCOLYSIS
ANDROGEN RESPONSE
INTERFERON GAMMA RESPONSE
CHOLESTEROL HOMEOSTASIS
ESTROGEN RESPONSE LATE
MITOTIC SPINDLE
PROTEIN SECRETION
TGF BETA SIGNALING
HEDGEHOG SIGNALING
ADIPOGENESIS
PI3K AKT MTOR SIGNALING
UV RESPONSE UP
PANCREAS BETA CELLS
BILE ACID METABOLISM
INFLAMMATORY RESPONSE
XENOBIOTIC METABOLISM
FATTY ACID METABOLISM
REACTIVE OXYGEN SPECIES PATHWAY
ALLOGRAFT REJECTION
PEROXISOME
WNT BETA CATENIN SIGNALING
SPERMATOGENESIS
DNA REPAIR
OXIDATIVE PHOSPHORYLATION
MTORC1 SIGNALING
UNFOLDED PROTEIN RESPONSE
G2M CHECKPOINT
E2F TARGETS
MYC TARGETS V2
MYC TARGETS V1
AN3CA
VOA1066
E
Z-Score
KRT8
ANKRD1
KRT16
MYBPH
DYNLT1
SERINC5
KRT80
FOSL2
PRKG1
FLJ42393
ZSCAN16-AS1
KRT75
SLC35E2B
MBIP
CD46
RIMS4
PCK1
MAPRE2
NUDT4
MRPL42P5
CFL1P1
KRT86
IFITM1
TBR1
PIKFYVE
PRKCD
TTC13
KPNA2
UBR7
XRCC2
LINC01159
RHOT2
SLFN11
DUOX1
KRT20
HSPD1
PRPSAP2
LDHB
PTBP1
COL26A1
DHX34
NTN1
RPL32
RRP15
SURF2
CSE1L
BCL2L2
PPRC1
PCP4L1
Par. -Dox +Dox (ARID1A) -Dox +Dox (ARID1A)
AN3CA VOA1066
F
Upregulated Genes
AN3CA VOA1066
768 419 906
OLFML1,COL12A1 SDPR,GAS2L2,IL7R TMC3,LNX1,PLCE1-AS1 SIGLEC1,INMT,HKDC1 WNT5A,PSG9 MFAP4,KL MEOX1
TRABD2B,WNT6 KRT17,FOLR1,KRT80 ROR1,PRICKLE1,L1CAM SOX13,PRKCA ARID1A,ACTN4
REN,TH,P2RY14 SHROOM4,CYS1,ZBTB7C LRFN2,ADAMTS2,CELP CHRNA4,SLA,MYBPHL PDLIM1,BTBD11 ARHGEF26 INSRR
Downregulated Genes
AN3CA VOA1066
253 21 216
SLC24A4,LPPR4 PRLR,INSM2,SOX8 FCGR2A,SERPINA3 AR,HIF1A-AS1,FCGR2B MIA,KCNG4,HEY1 RGS4,ST8SIA6 NKAIN1,LAG3
TERT NR4A2 GLI1 MYC DDN
KRT81,DENND2D DMP1,CYSLTR2,HNF4A ANXA10,SPRR2D SLC17A9,MYHAS RPRML,FLNC,ARL9 OLIG1,CHRNE LIF,MMRN2 GATA6
G
Quantitative TMT-MS AN3CA cells
n=125 n=430
– log10 Adj. P-value
log2 FC
+Dox (ARID1A)/-Dox
VOA1066 cells
n=28 n=307
H
AN3CA cells
CDH1
TONSL
CLDN4
BRCA2
Dox (ARID1A) :
ATAC
HA-ARID1A
SMARCA4
SS18
BRD9
GLTSCR1
H3K27ac
Chr16:68,721,595-68,872,228
Chr8:145,639,683-145,684,298
Chr7:73,229,323-73,253,161
Chr13:32,848,154-33,036,301
Replicate:
-Dox(Control) +Dox (ARID1A)

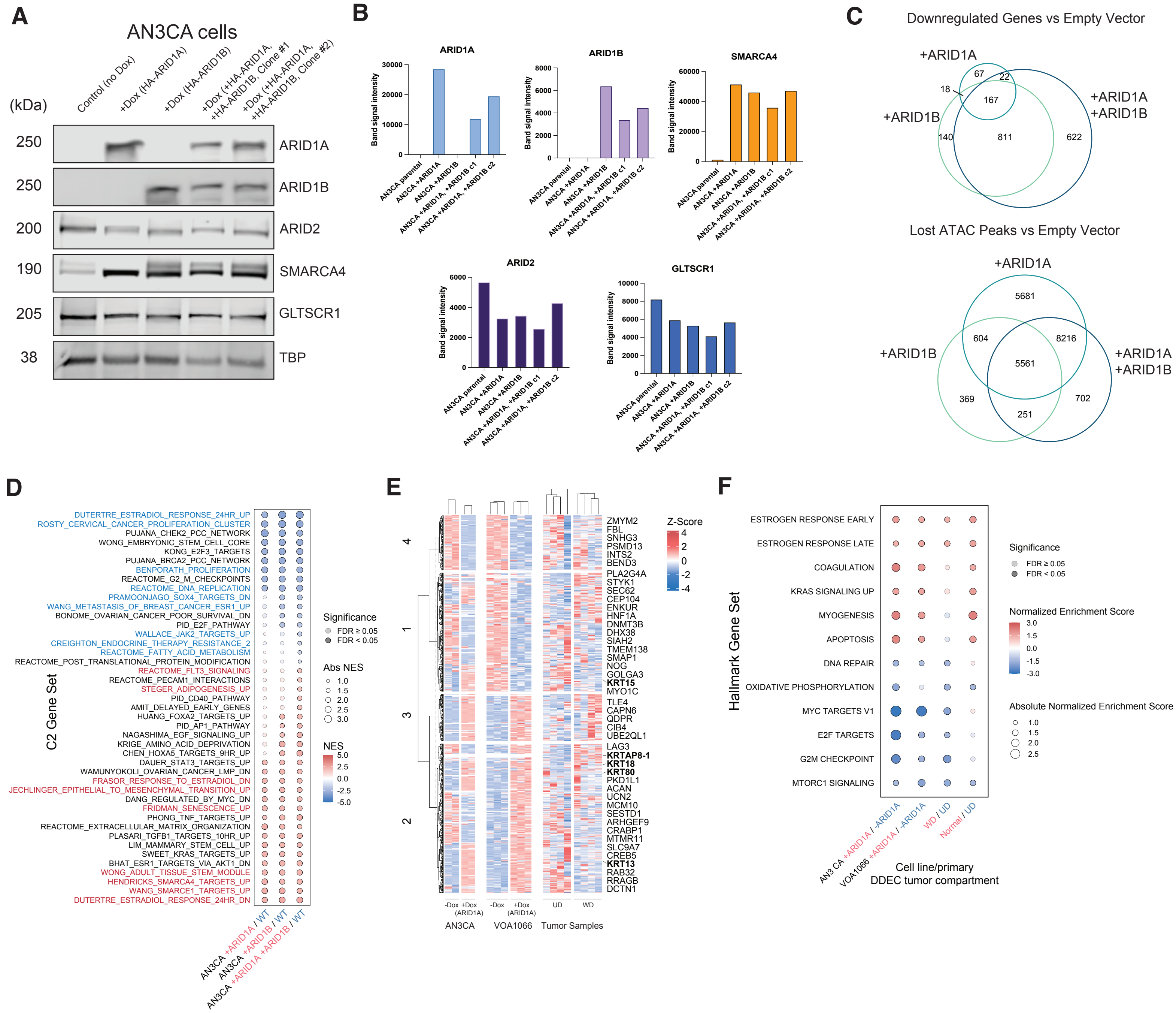
A
AN3CA cells
(kDa)
Control (no Dox)
+Dox (HA-ARID1A)
+Dox (HA-ARID1B)
+Dox (+HA-ARID1A, +HA-ARID1B, Clone #1
+Dox (+HA-ARID1A, +HA-ARID1B, Clone #2)
250 ARID1A
250 ARID1B
200 ARID2
190 SMARCA4
205 GLTSCR1
38 TBP
B
ARID1A
ARID1B
SMARCA4
ARID2
GLTSCR1
Band signal intensity
AN3CA parental
AN3CA +ARID1A
AN3CA +ARID1B
AN3CA +ARID1A, +ARID1B c1
AN3CA +ARID1A, +ARID1B c2
C
Downregulated Genes vs Empty Vector
+ARID1A
+ARID1B
+ARID1A +ARID1B
67
22
18
167
140
811
622
Lost ATAC Peaks vs Empty Vector
5681
604
8216
5561
369
702
251
D
C2 Gene Set
DUTERTRE_ESTRADIOL_RESPONSE_24HR_UP
ROSTY_CERVICAL_CANCER_PROLIFERATION_CLUSTER
PUJANA_CHEK2_PCC_NETWORK
WONG_EMBRYONIC_STEM_CELL_CORE
KONG_E2F3_TARGETS
PUJANA_BRCA2_PCC_NETWORK
BENPORATH_PROLIFERATION
REACTOME_G2_M_CHECKPOINTS
REACTOME_DNA_REPLICATION
PRAMOONJAGO_SOX4_TARGETS_DN
WANG_METASTASIS_OF_BREAST_CANCER_ESR1_UP
BONOME_OVARIAN_CANCER_POOR_SURVIVAL_DN
PID_E2F_PATHWAY
WALLACE_JAK2_TARGETS_UP
CREIGHTON_ENDOCRINE_THERAPY_RESISTANCE_2
REACTOME_FATTY_ACID_METABOLISM
REACTOME_POST_TRANSLATIONAL_PROTEIN_MODIFICATION
REACTOME_FLT3_SIGNALING
REACTOME_PECAM1_INTERACTIONS
STEGER_ADIPOGENESIS_UP
PID_CD40_PATHWAY
AMIT_DELAYED_EARLY_GENES
HUANG_FOXA2_TARGETS_UP
PID_AP1_PATHWAY
NAGASHIMA_EGF_SIGNALING_UP
KRIGE_AMINO_ACID_DEPRIVATION
CHEN_HOXA5_TARGETS_9HR_UP
DAUER_STAT3_TARGETS_UP
WAMUNYOKOLI_OVARIAN_CANCER_LMP_DN
FRASOR_RESPONSE_TO_ESTRADIOL_DN
JECHLINGER_EPITHELIAL_TO_MESENCHYMAL_TRANSITION_UP
DANG_REGULATED_BY_MYC_DN
FRIDMAN_SENESCENCE_UP
PHONG_TNF_TARGETS_UP
REACTOME_EXTRACELLULAR_MATRIX_ORGANIZATION
PLASARI_TGFB1_TARGETS_10HR_UP
LIM_MAMMARY_STEM_CELL_UP
SWEET_KRAS_TARGETS_UP
BHAT_ESR1_TARGETS_VIA_AKT1_DN
WONG_ADULT_TISSUE_STEM_MODULE
HENDRICKS_SMARCA4_TARGETS_UP
WANG_SMARCE1_TARGETS_UP
DUTERTRE_ESTRADIOL_RESPONSE_24HR_DN
Significance
FDR ≥ 0.05
FDR < 0.05
Abs NES
NES
AN3CA +ARID1A / WT
AN3CA +ARID1B / WT
AN3CA +ARID1A +ARID1B / WT
E
Z-Score
ZMYM2
FBL
SNHG3
PSMD13
INTS2
BEND3
PLA2G4A
STYK1
SEC62
ENKUR
CEP104
HNF1A
DNMT3B
DHX38
SIAH2
TMEM138
SMAP1
NOG
GOLGA3
KRT15
MYO1C
TLE4
CAPN6
QDPR
CIB4
UBE2QL1
LAG3
KRTAP8-1
KRT18
KRT80
PKD1L1
ACAN
UCN2
MCM10
SESTD1
ARHGEF9
CRABP1
MTMR11
SLC9A7
CREB5
KRT13
RAB32
RRAGB
DCTN1
-Dox
+Dox (ARID1A)
UD
WD
AN3CA
VOA1066
Tumor Samples
F
Hallmark Gene Set
ESTROGEN RESPONSE EARLY
ESTROGEN RESPONSE LATE
COAGULATION
KRAS SIGNALING UP
MYOGENESIS
APOPTOSIS
DNA REPAIR
OXIDATIVE PHOSPHORYLATION
MYC TARGETS V1
E2F TARGETS
G2M CHECKPOINT
MTORC1 SIGNALING
Significance
FDR ≥ 0.05
FDR < 0.05
Normalized Enrichment Score
Absolute Normalized Enrichment Score
AN3 CA +ARID1A / -ARID1A
VOA1066 +ARID1A / -ARID1A
WD / UD
Normal / UD
Cell line/primary
DDEC tumor compartment

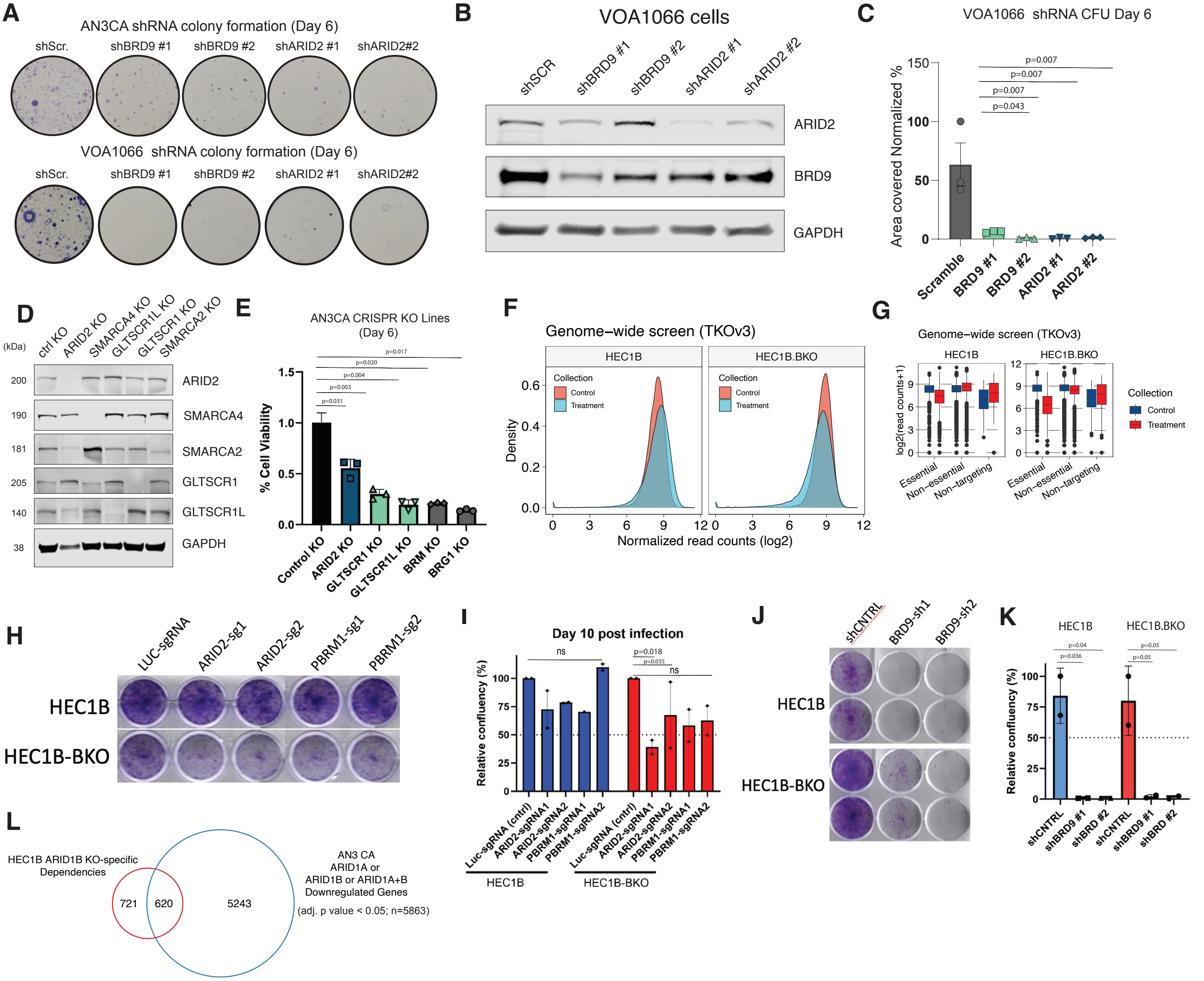
A
AN3CA shRNA colony formation (Day 6)
shScr. shBRD9 #1 shBRD9 #2 shARID2 #1 shARID2#2
VOA1066 shRNA colony formation (Day 6)
B
VOA1066 cells
shSCR shBRD9 #1 shBRD9 #2 shARID2 #1 shARID2 #2
ARID2
BRD9
GAPDH
C
VOA1066 shRNA CFU Day 6
Area covered Normalized %
p=0.007
p=0.043
Scramble BRD9 #1 BRD9 #2 ARID2 #1 ARID2 #2
D
ctrl KO ARID2 KO SMARCA4 KO GLTSCR1L KO GLTSCR1 KO SMARCA2 KO
(kDa)
ARID2 SMARCA4 SMARCA2 GLTSCR1 GLTSCR1L GAPDH
E
AN3CA CRISPR KO Lines (Day 6)
% Cell Viability
Control KO ARID2 KO GLTSCR1 KO GLTSCR1L KO BRM KO BRG1 KO
F
Genome−wide screen (TKOv3)
HEC1B HEC1B.BKO
Collection
Control
Treatment
Density
Normalized read counts (log2)
G
Genome−wide screen (TKOv3)
log2(read counts+1)
Essential Non−essential Non−targeting
H
LUC-sgRNA ARID2-sg1 ARID2-sg2 PBRM1-sg1 PBRM1-sg2
HEC1B
HEC1B-BKO
I
Day 10 post infection
Relative confluency (%)
ns
p=0.018
p=0.035
Luc-sgRNA (cntrl) ARID2-sgRNA1 ARID2-sgRNA2 PBRM1-sgRNA1 PBRM1-sgRNA2
J
shCNTRL BRD9-sh1 BRD9-sh2
K
p=0.04
p=0.036
p=0.05
shCNTRL shBRD9 #1 shBRD #2
L
HEC1B ARID1B KO-specific Dependencies
721 620 5243
AN3 CA ARID1A or ARID1B or ARID1A+B Downregulated Genes
(adj. p value < 0.05; n=5863)

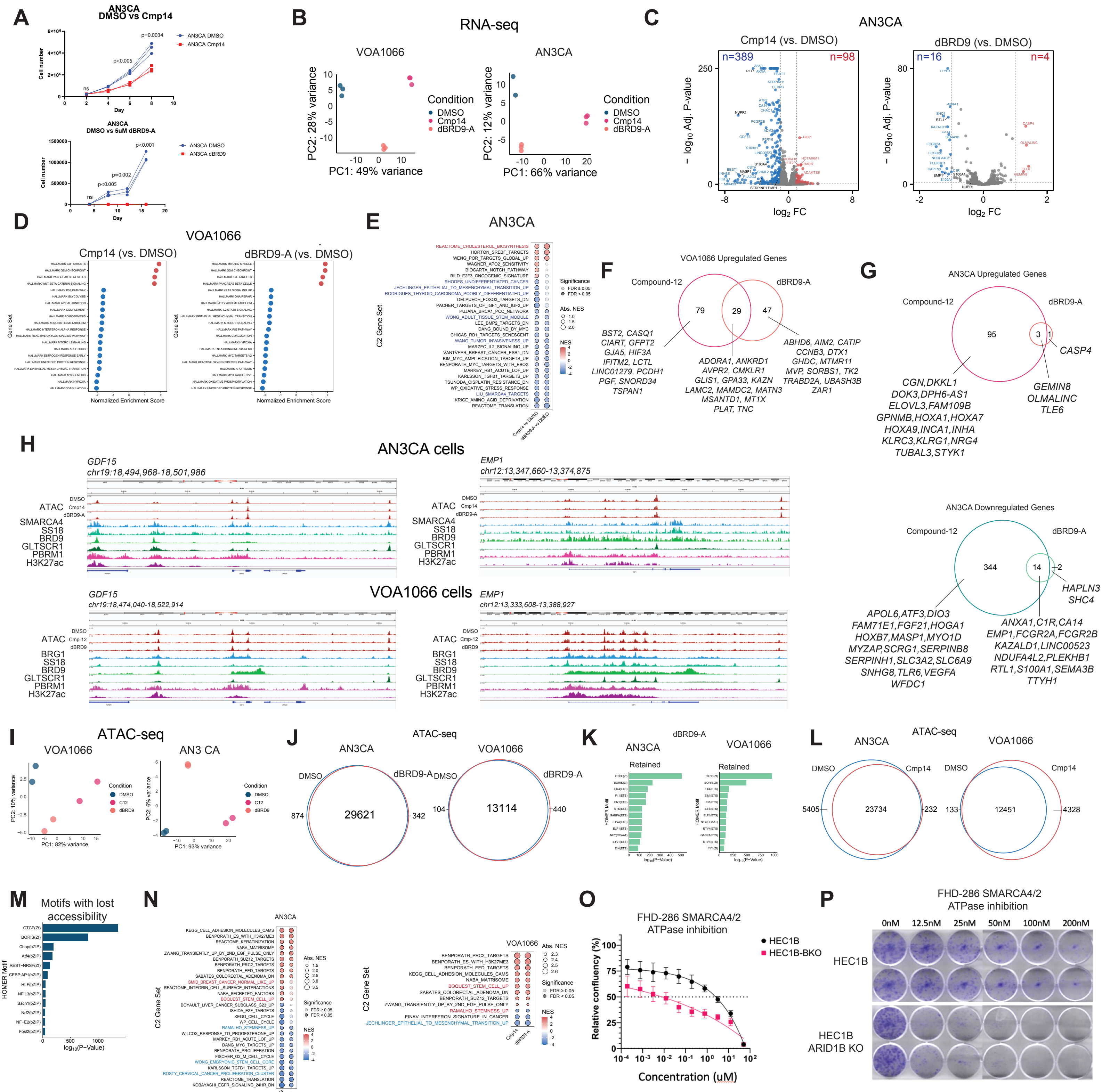
A
AN3CA DMSO vs Cmp14
AN3CA DMSO vs 5uM dBRD9-A
B
RNA-seq
VOA1066
AN3CA
Condition
DMSO
Cmp14
dBRD9-A
PC1: 49% variance
PC2: 28% variance
PC1: 66% variance
PC2: 12% variance
C
AN3CA
Cmp14 (vs. DMSO)
dBRD9 (vs. DMSO)
n=389
n=98
n=16
n=4
D
VOA1066
Cmp14 (vs. DMSO)
dBRD9-A (vs. DMSO)
Normalized Enrichment Score
E
AN3CA
C2 Gene Set
F
VOA1066 Upregulated Genes
Compound-12
dBRD9-A
79
29
47
BST2, CASQ1 CIART, GFPT2 GJA5, HIF3A IFITM2, LCTL LINC01279, PCDH1 PGF, SNORD34 TSPAN1
ADORA1, ANKRD1 AVPR2, CMKLR1 GLIS1, GPA33, KAZN LAMC2, MAMDC2, MATN3 MSANTD1, MT1X PLAT, TNC
ABHD6, AIM2, CATIP CCNB3, DTX1 GHDC, MTMR11 MVP, SORBS1, TK2 TRABD2A, UBASH3B ZAR1
G
AN3CA Upregulated Genes
Compound-12
dBRD9-A
95
3
1
CASP4
CGN,DKKL1 DOK3,DPH6-AS1 ELOVL3,FAM109B GPNMB,HOXA1,HOXA7 HOXA9,INCA1,INHA KLRC3,KLRG1,NRG4 TUBAL3,STYK1
GEMIN8 OLMALINC TLE6
AN3CA Downregulated Genes
Compound-12
dBRD9-A
344
14
2
HAPLN3 SHC4
APOL6,ATF3,DIO3 FAM71E1,FGF21,HOGA1 HOXB7,MASP1,MYO1D MYZAP,SCRG1,SERPINB8 SERPINH1,SLC3A2,SLC6A9 SNHG8,TLR6,VEGFA WFDC1
ANXA1,C1R,CA14 EMP1,FCGR2A,FCGR2B KAZALD1,LINC00523 NDUFA4L2,PLEKHB1 RTL1,S100A1,SEMA3B TTYH1
H
AN3CA cells
GDF15 chr19:18,494,968-18,501,986
EMP1 chr12:13,347,660-13,374,875
ATAC
SMARCA4
SS18
BRD9
GLTSCR1
PBRM1
H3K27ac
VOA1066 cells
GDF15 chr19:18,474,040-18,522,914
EMP1 chr12:13,333,608-13,388,927
BRG1
I
ATAC-seq
VOA1066
AN3 CA
J
ATAC-seq
AN3CA
VOA1066
DMSO
dBRD9-A
874
29621
342
104
13114
440
K
dBRD9-A
AN3CA
VOA1066
Retained
L
ATAC-seq
AN3CA
VOA1066
DMSO
Cmp14
5405
23734
232
133
12451
4328
M
Motifs with lost accessibility
N
AN3CA
VOA1066
C2 Gene Set
O
FHD-286 SMARCA4/2 ATPase inhibition
HEC1B
HEC1B-BKO
Relative confluency (%)
Concentration (uM)
P
FHD-286 SMARCA4/2 ATPase inhibition
0nM
12.5nM
25nM
50nM
100nM
200nM
HEC1B
HEC1B ARID1B KO

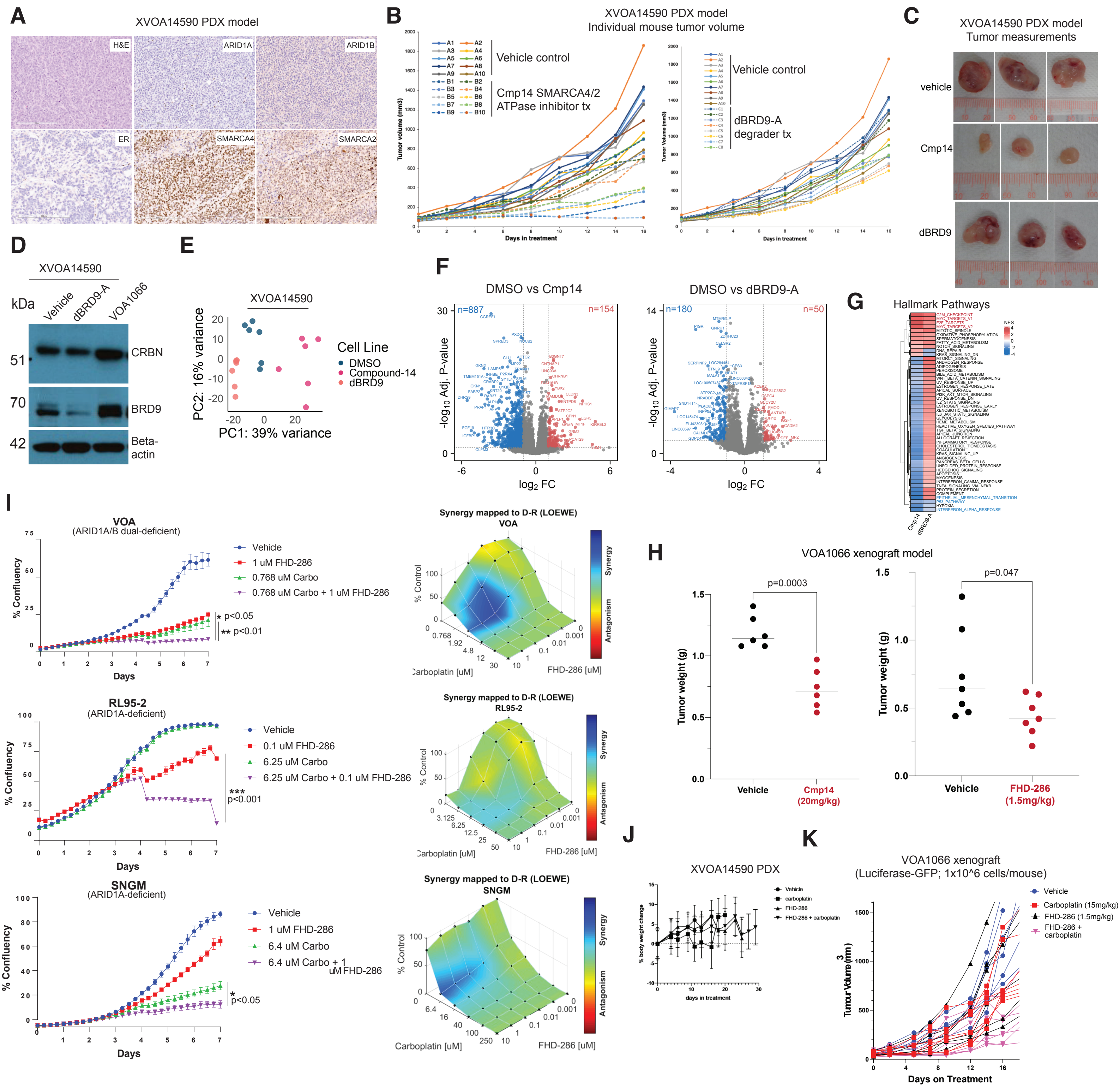
A
XVOA14590 PDX model
H&E
ARID1A
ARID1B
ER
SMARCA4
SMARCA2
B
XVOA14590 PDX model
Individual mouse tumor volume
Vehicle control
Cmp14 SMARCA4/2 ATPase inhibitor tx
Tumor volume (mm3)
Days in treatment
Vehicle control
dBRD9-A degrader tx
Tumor Volume (mm3)
Days in treatment
C
XVOA14590 PDX model
Tumor measurements
vehicle
Cmp14
dBRD9
D
XVOA14590
kDa
Vehicle
dBRD9-A
VOA1066
51
CRBN
70
BRD9
42
Beta-actin
E
XVOA14590
PC2: 16% variance
PC1: 39% variance
Cell Line
DMSO
Compound-14
dBRD9
F
DMSO vs Cmp14
n=887
n=154
-log10 Adj. P-value
log2 FC
DMSO vs dBRD9-A
n=180
n=50
-log10 Adj. P-value
log2 FC
G
Hallmark Pathways
NES
Cmp14
dBRD9-A
I
VOA
(ARID1A/B dual-deficient)
% Confluency
Days
Vehicle
1 uM FHD-286
0.768 uM Carbo
0.768 uM Carbo + 1 uM FHD-286
* p<0.05
** p<0.01
Synergy mapped to D-R (LOEWE)
VOA
% Control
Carboplatin [uM]
FHD-286 [uM]
Synergy
Antagonism
RL95-2
(ARID1A-deficient)
% Confluency
Days
Vehicle
0.1 uM FHD-286
6.25 uM Carbo
6.25 uM Carbo + 0.1 uM FHD-286
***
p<0.001
Synergy mapped to D-R (LOEWE)
RL95-2
% Control
Carboplatin [uM]
FHD-286 [uM]
Synergy
Antagonism
SNGM
(ARID1A-deficient)
% Confluency
Days
Vehicle
1 uM FHD-286
6.4 uM Carbo
6.4 uM Carbo + 1 uM FHD-286
*
p<0.05
Synergy mapped to D-R (LOEWE)
SNGM
% Control
Carboplatin [uM]
FHD-286 [uM]
Synergy
Antagonism
H
VOA1066 xenograft model
p=0.0003
Tumor weight (g)
Vehicle
Cmp14 (20mg/kg)
p=0.047
Tumor weight (g)
Vehicle
FHD-286 (1.5mg/kg)
J
XVOA14590 PDX
Vehicle
carboplatin
FHD-286
FHD-286 + carboplatin
% body weight change
days in treatment
K
VOA1066 xenograft
(Luciferase-GFP; 1x10^6 cells/mouse)
Vehicle
Carboplatin (15mg/kg)
FHD-286 (1.5mg/kg)
FHD-286 + carboplatin
Tumour Volume (mm3)
Days on Treatment
